# Typhoon-induced Lammas growth promotes the non-dormant life-cycle of the Great Orange Tip butterfly *Hebomoia glaucippe*

**DOI:** 10.1101/2023.05.09.539972

**Authors:** Kota Ogawa, Wataru Nakamizo, Fukashi Ishiwata, Yu Matsuura, Akiko Satake

**Affiliations:** Biosystematics Laboratory, Faculty of Social and Cultural Studies, Kyushu University, 744 Motooka, Nishi-ku, Fukuoka, 819-0395, Japan; Insect Science and Creative Entomology Center, Kyushu University, 744 Motooka, Nishi-ku, Fukuoka, 819-0395, Japan; Department of Biology, Faculty of Science, Kyushu University, 744 Motooka, Nishi-ku, Fukuoka, 819-0395, Japan; Graduate School of Integrated Science for Global Society, Kyushu University, 744 Motooka, Nishi-ku, Fukuoka 819−0395 Japan; 1140-4 Kurio, Yakushima-cho, Kumage-gun, Kagoshima, 891-4409, Japan; Tropical Biosphere Research Center, University of the Ryukyus, Nishihara, Okinawa, 903-0213, Japan

**Keywords:** Convergent Cross Mapping (CCM), Empirical Dynamic Modeling (EDM), life-cycle polyphenism, overwintering strategy, phenology, population dynamics, seasonal adaptation

## Abstract

Dormancy is a significant adaptation that enables organisms to overcome unfavorable seasons for survival and reproduction. Therefore, understanding the origin of dormancy is critical to comprehend the adaptation of tropical organisms to cold climates at high latitudes, limiting factors of their distribution. The great orange-tip butterfly *Hebomoia glaucippe* and its subspecies in East Asia exhibit various dormancy features and thus offer a suitable model for exploring seasonal adaptations in animals. Here, we investigated the dormancy of three subspecies of *H. glaucippe* in Japan: *shirozui*, *liukiuensis*, and *cincia*. Our rearing experiments indicate that *shirozui* and *liukiuensis* enter dormancy during the pupal stage under low temperatures and short-day conditions. Conversely, ssp. *cincia* does not exhibit dormancy even under similar conditions. Although *Crateva religiosa*, the only host plant in Japan, is deciduous, our field survey revealed that typhoon disturbances in autumn induce Lammas growth, and the following secondary shoots are used as a food source during winter in Yaeyama Islands, where ssp. *cincia* is distributed. Analyzing 10-year population dynamics and meteorological data, we demonstrated the seasonal occurrence of ssp. *liukiuensis* in Okinawa Island but not that of ssp. *cincia* in Ishigaki Island. By employing a non-linear time series causal inference framework, we discovered that temperature was causally related to butterfly occurrence on Okinawa Island, whereas maximum wind speed and precipitation had a causal relationship to butterfly occurrence on Ishigaki Island. The optimal time lag for both environmental factors that affect the population was approximately 60 days on Ishigaki Island, corresponding to the time from egg to adult in *H. glaucippe*. Collectively, these environmental factors are likely to determine the dynamics of the next generation of the Ishigaki population. Our findings suggest that frequent typhoons disturb seasonal defoliation cycles of the host plant and accompany Lammas growth in the Ryukyu Islands, reducing the selection pressure in the adaptation of *H. glaucippe* for cyclically-changing seasons, leading to the absence of dormancy in ssp. *cincia*.

**Significance:** Many organisms have evolved adaptive life histories to respond to seasonal and periodic environmental fluctuations. On the other hand, random environmental disturbances such as hurricanes and wildfires often occur on Earth, and these unpredictable events are likely to impact the life histories of organisms. Through comprehensive investigation using long-term time-series data, we proposed that typhoon-generated environmental disturbances may counteract the acquisition of dormancy by *Hebomoia glaucippe* in Japan. Our study highlights the significant impact of environmental disturbances on population dynamics and emphasizes that these frequent disturbances may obscure the influence of seasonal environmental changes, thereby promoting multifarious life-cycle evolution.

## Introduction

Organisms have developed various strategies to adapt to cyclical seasonal changes. Especially for insects with short lifespans and small body sizes, surviving unsuitable seasons such as winter and the dry season is a critical challenge (Denlinger, 2022). Diapause, an adaptive physiological state controlled by the endocrine system, allows insects to tolerate severe conditions such as cold and drought, playing a crucial role in seasonal adaptation and life history evolution (Koštál 2006, Gill et al., 2017; Tougeron, 2019; Denlinger, 2022; Goto, 2022). Since inappropriately timed dormancy leads to missed opportunities for growth and mating, the environmental signals (environmental cues) used to induce and break dormancy are extremely important. Photoperiod (day length) serves as a reliable signal for insects to detect seasonal changes due to its consistent features that vary depending on season and latitude while exhibiting low interannual fluctuation (Gill et al., 2017; Tougeron, 2019; Denlinger, 2022). Moreover, environmental factors fluctuate with changes in day length, such as temperature and humidity, often modifying the photoperiodic response. In particular, the temperature is the second most crucial factor after day length; the critical day length for inducing dormant pupae increases as temperature decreases in some insects (e.g., Kato, 2000). In tropical and subtropical regions with low latitudes, environmental factors such as temperature, humidity, and food availability play more critical roles as dormancy-inducing signals, given the less variable seasonal photoperiod in these regions (Gill et al., 2019; Tougeron, 2019; Denlinger, 2022).

Determining biotic and abiotic factors driving environmentally-induced dormancy is crucial for understanding the seasonal adaptation and the life-cycle evolution of insects. The Great Orange Tip butterfly *Hebomoia glaucippe* is a suitable model for exploring transitional changes of life history traits since it exhibits various types of pupal dormancy among the populations (Ogawa 2019). Although *Hebomoia glaucippe* is widely distributed throughout Southeast Asia and the Indian subcontinent, the northernmost distribution is along the Ryukyu Islands in the Japanese archipelago (Morishita 1973, Samusawa, 2021, Fig. 1a). *H. glaucippe* is not dormant in its native tropical habitat, but in subtropical Hong Kong, where the nominotypical subspecies *glaucippe* is distributed, its life cycle (dormant/non-dormant) varies based on the status of food plants (Bascombe et al., 1999). Specifically, when the evergreen *Capparis cantoniensis* is used as a food plant, the larvae continue to feed and grow throughout the winter and develop into non-dormant pupae. In contrast, when the deciduous *Crateva religiosa* is used as a food plant, the larvae become dormant pupae before leaves fall and overwinter as such. On the other hand, short-day conditions can induce dormant pupae irrespective of the food plant in the northernmost population in Kyushu, Japan (Yata 1977). Accordingly, despite *Hebomoia glaucippe* employing food plant-dependent pupal dormancy in the subtropical zone, daylength-dependent dormancy was acquired during their northern expansion into the Ryukyu Islands via Taiwan (Ogawa 2019).

**Figure 1.**
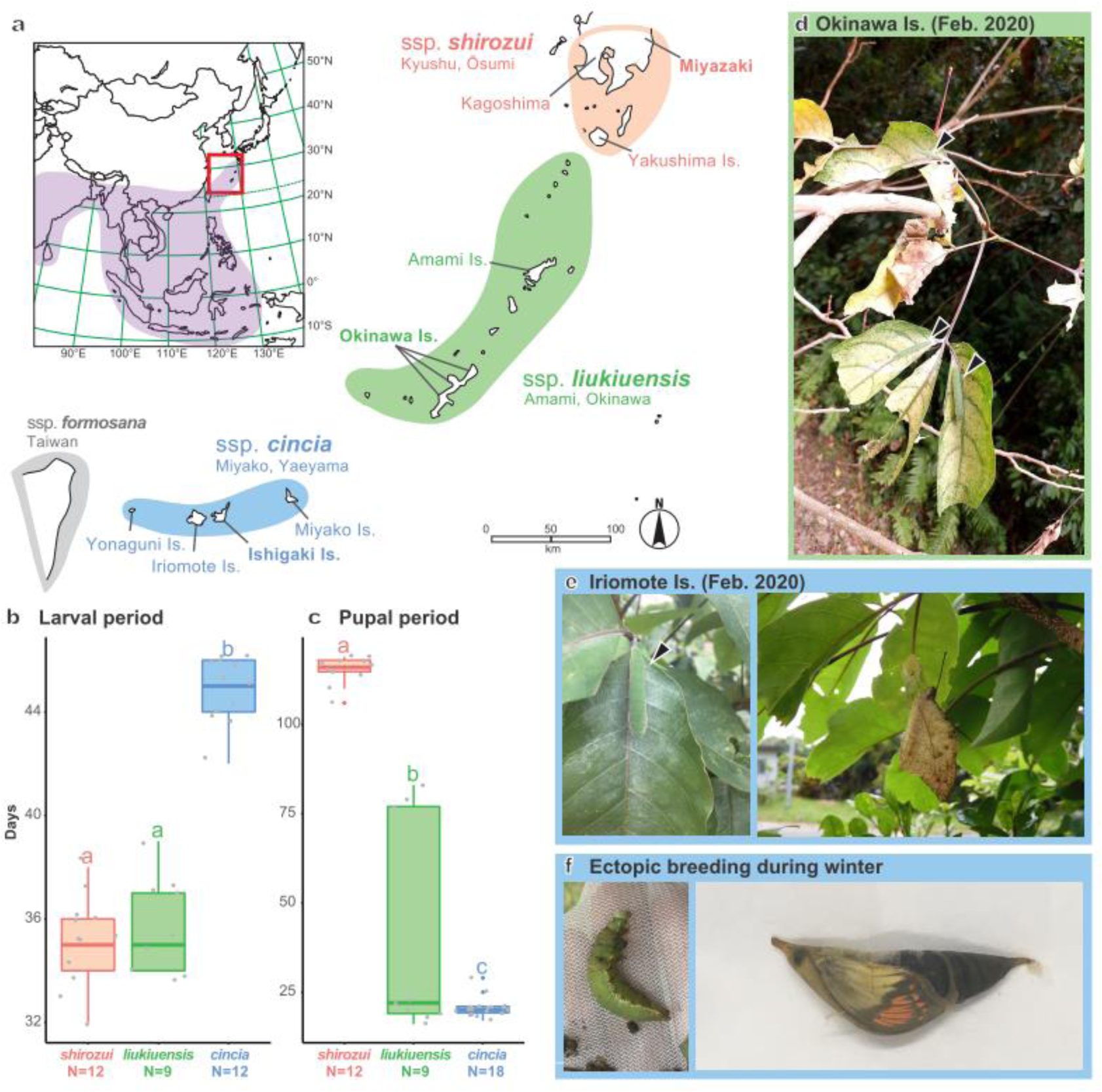
Distribution and overwintering status of *Hebomoia glaucippe*. **(a)** Distribution map. The violet shadow and the red box in the inset indicate the global distribution and the magnified area between Japan and Taiwan. The orange, green, and blue shadows on the magnified map indicate the distribution of subspecies *shirozui*, *liukiuensis*, and *cincia*, respectively. **(b, c)** Larval and pupal periods of each subspecies when reared at 12L12D, 20°C. Different letters indicate significant differences between groups (one-way ANOVA followed by a post-hoc Tukey HSD test, P<0.05). **(d, e)** The overwintering situation in the fields. In the subspecies *liukiuensis*, due to winter defoliation of hosts, it is difficult to survive the winter as a larva (d, supplemental figure S2). In the subspecies *cincia*, however, there are many hosts with leaves even in winter, and the larval and adult activity can be observed annually (e, supplemental figure S3). Arrowheads indicate larvae on leaves. **(f)** Dead larva and pupa during molting in the ectopic overwintering experiments. In Fukuoka City (33°35’58N 130°13’22E), neither larvae nor pupae die directly due to low temperatures; nonetheless, they cannot complete the molt before their external epidermis sclerotizes (see supplemental figure S4).

While some taxonomic issues remain unresolved, *H. glaucippe* comprises more than 40 described subspecies, with closely related subspecies grouped together to form several subspecies groups (Morishita, 1973; Treadaway & Schröder, 2008; Samusawa, 2021). In Japan, three subspecies, ssp. *shirozui* Kurosawa & Omoto, 1955, ssp. *liukiuensis* Fruhstorfer, 1898, and ssp. *cincia* Fruhstorfer, 1910, are found from Kyushu to Yaku Island, Amami to Okinawa Island, and Miyako to Yonaguni Island, respectively (Fig. 1a). These three subspecies belong to the *liukiuensis* group and are often collectively regarded as ssp. *liukiuensis* (reviewed in Samusawa, 2021). While pupal diapause induced by short-day conditions has been confirmed in the Kagoshima population of ssp. *shirozui* (Yata, 1977), it remains unknown whether other subspecies distributed in Japan exhibit the same dormancy. Additionally, in Japan, the deciduous host tree *Crateva religiosa* is the only host plant for *H. glaucippe* (Higa & Nagamine, 2019), in contrast to Hong Kong and Taiwan, where evergreen hosts are also distributed (Bascombe et al., 1999;). Therefore, studying the overwintering strategy and dormancy of *Hebomoia glaucippe* in the Ryukyu Archipelago will provide valuable insight into the dormancy-promoing life-cycle alternation.

The specific objectives of our study are to clarify dormancy variations among the *H. glaucippe* subspecies from three remote areas, the environmental factors involved in this process, and the impact of environmental parameters on population dynamics and subsequent life-cycle alternation. To accomplish the above, we conducted investigations of dormancy in each population in captivity and examined overwintering conditions in the field. Our field observations serendipitously found that typhoon disturbances affect *H. glaucippe* emergence, and therefore, we statistically examined how typhoons and the unique climate of the Ryukyu Islands affect the population dynamics of *H. glaucippe*. Specifically, we used the continual 10 years of *Hebomoia glaucippe* occurrence data and meteorological data to evaluate the long-term effects of environmental factors on population dynamics. Based on these ecological and mathematical findings, we will discuss how selection pressure is altered and how dormancy responsiveness is affected by randomly occurring weather events that override cyclical and seasonal changes.

## Results

### Shortday response of *Hebomoia glaucippe* in Japan

To compare the short-day response and diapause-related life-history traits, we reared each subspecies of *H. glaucippe* under low-temperature and short-day conditions (12L12D20°C) (Figs. 1b,c). Notably, vernalization treatment was unnecessary for the emergence of the diapause pupae, and thus, all pupae were cultured in identical conditions as larvae. The larval period was about 30% longer in ssp. *cincia* (44.8 ± 1.27 days [mean ± S.D.], n = 12) than in ssp. *shirozui* (35.1 ± 1.68 days [mean ± S.D.], n = 12) and *liukiuensis* (35.8 ± 1.79 days [mean ± S.D.], n = 9), significantly (one-way ANOVA followed by a post-hoc Tukey HSD test, P<0.05; Fig. 1b).

As discerning between diapause and non-diapause pupae of this species is unfeasible through external morphology (Yata, 1977), we evaluated the expression of diapause by analyzing the duration of the pupal period. Previous studies showed that the pupal period under long-day conditions is less than 20 days (Yata, 1977, Bascombe et al., 1999). Accordingly, we have operationally defined diapausing pupae as those with a pupal period exceeding 40 days, which is more than twice the duration of those raised under long-day conditions. The pupal periods of ssp. *shirozui* (115.3 ± 3.86 days [mean ± S.D.], n = 12), *liukiuensis* (39.7 ± 30.10 days [mean ± S.D.], n = 9), and *cincia* (20.7 ± 2.66 days [mean ± S.D.], n = 18) differed significantly (one-way ANOVA followed by a post-hoc Tukey HSD test, P<0.05; Fig. 1c). In ssp. *cincia*, only individuals with short pupal periods of about three weeks appeared (thus, all were non-diapause), whereas there were diapause pupae that took more than two months to emerge in the other two subspecies. While diapause occurred in all pupae of ssp. *shirozui*, which had the longest pupal period, both pupae were induced in ssp. *liukiuensis*, with 33% (3/9) diapause pupae. The pupal period of diapause pupae of ssp. *liukiuensis* (approx. 80 days) was shorter than that of ssp. *shirozui*, suggesting a regional cline of diapause duration.

### The overwintering condition of *Hebomoia glaucippe* in the fields

Although the rearing experiments have shown that all individuals of ssp. *cincia* and some of ssp. *liukiuensis* develop into non-diapause pupae even under short-day conditions, *Crateva religiosa* is deciduous and the only host tree of this butterfly species in Japan. Therefore, we investigated how these larvae or pupae overwinter in the field and whether non-diapause pupae can survive *in situ*. From 2017 to 2023, field observations were conducted in Miyazaki Prefecture (Nichinan City), Kagoshima Prefecture (Ibusuki City), Yakushima Island, Amami-Oshima Island, Okinawa Island, Miyako Islands, Ishigaki Island, Iriomote Island, and Yonaguni Island (Fig. 1a).

Most *Crateva religiosa* trees on Kyushu and Yakushima Island, where ssp. *shirozui* is distributed, shed their leaves during winter (Fig. S1). Moreover, many diapausing pupae were found on the undersides of the leaves and branches of plants nearby host trees before the defoliation occurred (these pupae were identified as diapausing pupae after being reared indoors, exhibiting a pupal period of 40 days or more; Fig. S1). Similarly, on Okinawa Island, where ssp. *liukiuensis* is distributed, most *C. religiosa* trees shed their leaves during winter (Fig. S2). Larvae that had matured enough before the defoliation abandoned their hosts and pupated on the undersides of the leaves and branches of nearby evergreen trees. Individuals that remained larvae throughout the winter would forage for food, leading many larvae to gather on the few remaining leaves (Fig. 1d, S2). Given this situation, the overwintering success rate of non-diapause individuals and the reproductive success rate of adults that emerged during the winter might be limited, at least for the years we observed.

In the Yaeyama Islands (Miyako Islands, Ishigaki Island, Iriomote Island, and Yonaguni Island), however, *C. religiosa* trees that retain their leaves during winter were frequently found, resulting in an asynchrony of leaf phenology among trees even within the same island (Fig. S3). The asynchrony of leaf phenology among host plants provide fresh leaves to *Hebomoia glaucippe* throughout the year on these islands; thereby, all developmental stages of *H. glaucippe* were observed even in winter (Fig. 1e, S3). It is, therefore, evident that ssp. *cincia* can endure the winter without undergoing diapause by relying on hosts that retain their leaves. However, it is noteworthy ssp. *cincia* is unable to enter diapause and cease development, causing it to molt inadequately and die in severe winter conditions (Fig. 1f, S4).

### Typhoon-induced disturbances and Lammas growth of *C. religiosa*

Then, we investigated the cause of the asynchronous phenology of *C. religiosa* on the Yaeyama Islands, where the ssp. *cincia* overwinter in a non-diapause mode. For year-round observations, we transplanted three *C. religiosa* trees to our private garden on Iriomote Island and used them as specimen trees. In addition to these specimens, we observed more than ten *C. religiosa* trees on Iriomote Island, depending on the prevailing circumstances, from April 2019 to March 2021. Our observation revealed that autumn typhoons significantly impact the phenology of *C. religiosa*. The leaves of *C. religiosa* are observed to become yellow gradually, with old leaves tending to persist for an extended period (Fig. S5). The *C. religiosa* leaves drop simultaneously due to typhoon-induced disturbances (Fig. 2a). Subsequently, Lammas growth, characterized by the growth of secondary shoots and leaves to compensate for leaf damage, occurs and attracts female *H. glaucippe*, which rapidly lays eggs on the Lammas leaves (Fig. 2b,c, S6).

**Figure 2.**
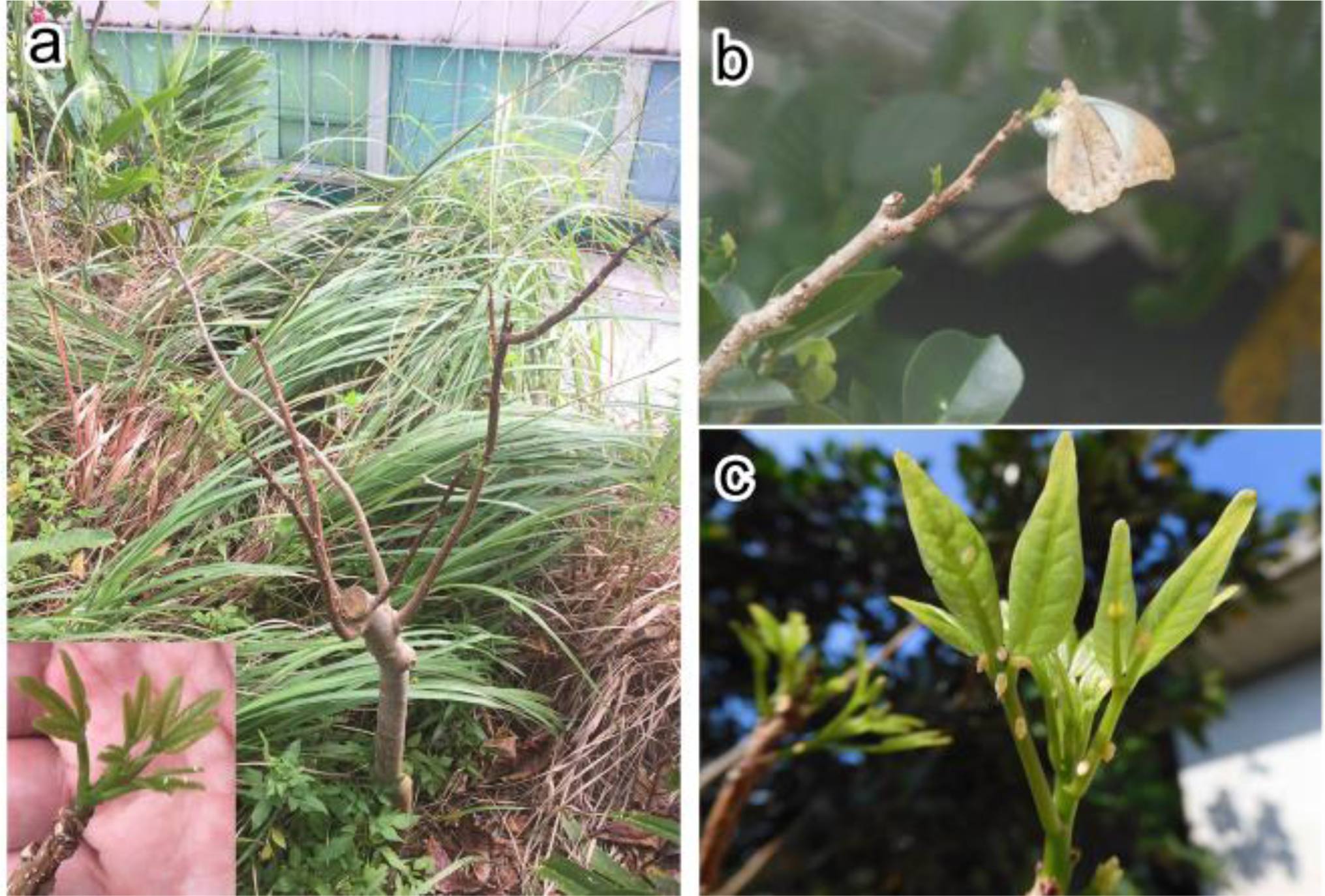
The typhoon disturbance induced Lammas growth of *Crateva religiosa* and ferocious oviposition on the Lammas shoots by *Hebomoia glaucippe*. **(a)** *C. religiosa* with the Lammas shoots. The Lammas shoot is shown in the inset. The typhoon defoliated all the old leaves, and a while later, new shoots sprouted. The leaves of typhoon-induced Lammas shoots do not fall off during winter; in other words, the Lammas growth induced trees will have healthy leaves during winter. **(b)** *H. glaucippe* ovipositing on the Lammas shoot. **(c)** Numerous eggs laid on the Lammas shoots.

Since the Lammas leaves sprouted in autumn do not defoliate over the winter, Lammas growth considerably influences the winter phenology of *C. religiosa* and the non-dormant life-cycle of ssp. *cincia*. However, the induction of the Lammas growth is contingent upon several variables, including the growing environment of *C. religiosa* and the course and intensity of the typhoon that can affect defoliation. Temperature also plays a role in determining the sprouting and subsequent shoot growth after the defoliation. Moreover, our limited field observations were ambiguous to ascertain and correlate the ubiquity of typhoon-induced disturbances, the consequent Lammas growth, and their effects on butterfly population dynamics and life cycles. Thus, we deemed it necessary to conduct a comprehensive analysis of long-term butterfly population dynamics along with meteorological data.

### Verification of the association between typhoon and butterfly population dynamics using long-term data

To verify the impact of typhoon disturbance and subsequent Lammas growth on the population dynamics of *H. glaucippe*, we compiled local collection records of adult individuals published by Japanese butterfly enthusiasts. We obtained data for Miyazaki (ssp. *shirozui*) over four years (2015-2018) and for Okinawa Island (ssp. *liukiuensis*) and Ishigaki Island (ssp. *cincia*) over ten years (2010-2019) (see **Materials & Methods** and **Supplemental Table** for details). First, we compared the population dynamics in the three regions. While there were only a few records in Miyazaki and Okinawa Island during the winter, adult butterflies were recorded throughout the year on Ishigaki Island (Fig. 3a). The autocorrelation analysis revealed a significant autocorrelation at approximately 180 and 360-day time lags on Okinawa Island, indicating a semi-annual and annual cycle of adult emergence (Fig. S7a). In contrast, on Ishigaki Island, no periodicity in occurrence was observed (Fig. S7b). The Miyazaki population datasets, including only 4-year averages, could not be used for the ACF analysis. The yearly emergence patterns in three islands were clustered into four groups (#1-4; Fig. 3b). Clusters #1 and #2, comprising data from Ishigaki Island, were characterized by the presence of adult individuals during the winter season and minimal seasonal fluctuations in occurrence. While the primary factor separating clusters #1 and #2 is unclear, cluster #2 showed an increase in occurrences in late March, followed by a decline. Clusters #3 and #4 both had limited adult records during winter, but cluster #3 was dominated by a high proportion of spring records (April-June), while cluster #4 was dominated by a high proportion of autumn records (Oct-Nov). Cluster #3 comprised only data from Okinawa, while cluster #4 included data from Ishigaki for two years in addition to Miyazaki and Okinawa.

**Figure 3.**
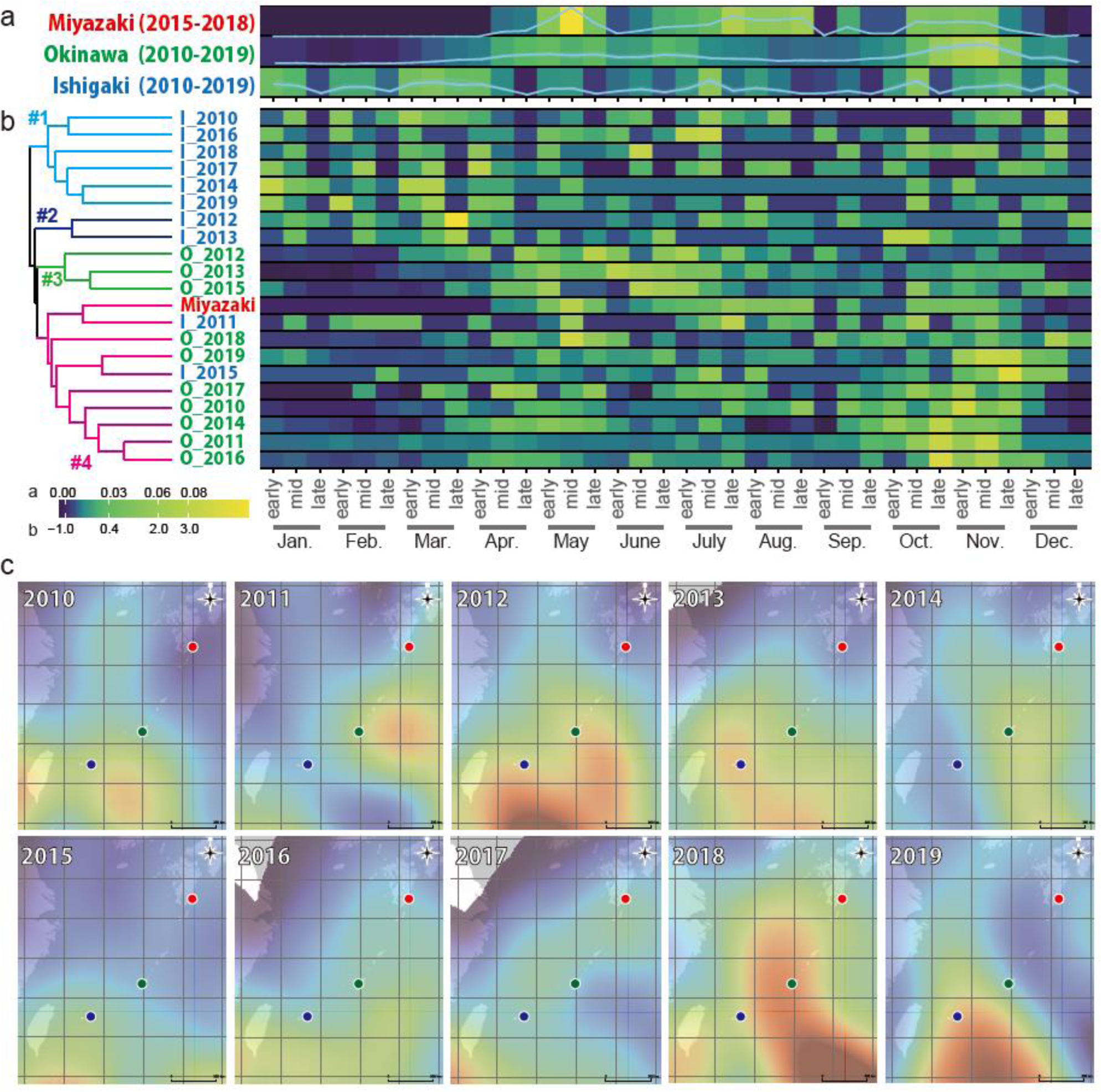
The population dynamics of *Hebomoia glaucippe* and the estimated typhoon impacts. **(a)** A period-averaged population dynamics heatmap for each population. **(b)** Hierarchical clustering heatmap of the population dynamics for each year. Noting that data for each year were unavailable for the Miyazaki population, the period average was used for the clustering analysis. The four clusters are indicated with numbers (#1-4) and branch colors. **(c)** Location-based estimated typhoon impact. Heatmaps were produced based on the typhoon center position information captured every six hours. Areas with a large number of center plots, i.e., areas considered to have been heavily impacted, are shown in red, while areas with a small number of center plots, i.e., areas considered to have been less disturbed, are depicted in blue. The red, green, and blue dots indicate the locations of Miyazaki (Nichinan City), Okinawa Island (Naha city), and Ishigaki Island, respectively.

Given that specific temperature thresholds are required for plant sprouting and budding (Horvath et al., 2003; Kim et al., 2009), we also investigated the association between typhoon attacks and temperatures in the three areas (Fig. 4). The number of typhoons approaching within 300 km of the meteorological observatory does not differ significantly among the three areas, and typhoons frequently occur between July to October (Fig. 4b). While there is no noticeable difference in temperatures during the typhoon season among the three areas, temperatures in Miyazaki decrease rapidly after October (Fig. 4a). Conversely, Okinawa and Ishigaki exhibit little difference in temperature until December, while Okinawa experiences a decrease in temperature in January and February of the new year (Fig. 4a). These data suggest that Lammas growth is more likely to occur in the southern region.

**Figure 4.**
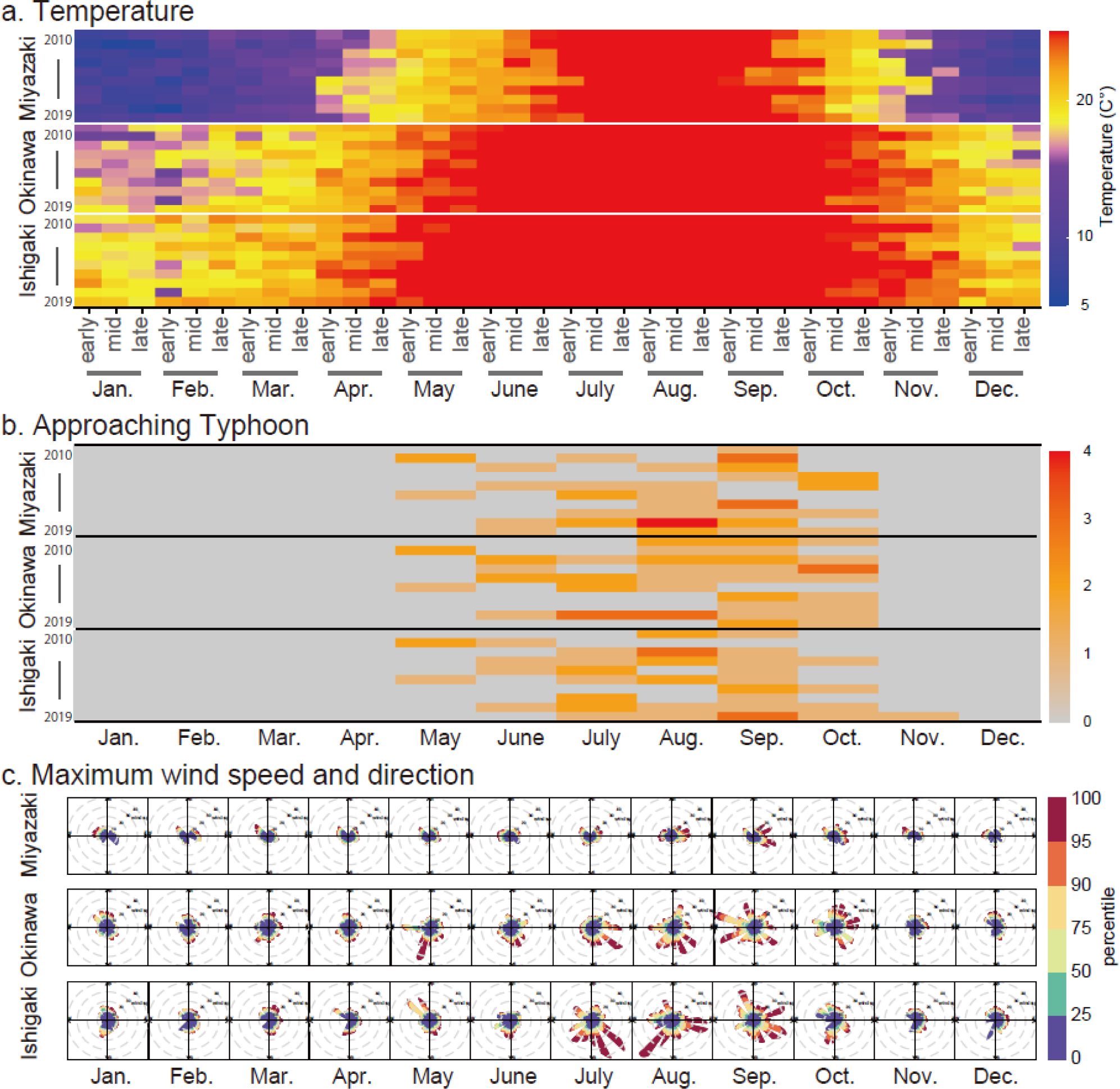
Comparison of environmental parameters for Miyazaki, Okinawa Island, and Ishigaki Island for 2010-2019. **(a)** Heatmap of the average temperature. **(b)** Heatmap of the number of the approaching typhoon. Approaching typhoons are defined as typhoons that reach within 300 km of the meteorological stations. **(c)** The percentile rose of the maximum wind speed and wind direction. The gray circles indicate wind speed scales of 10, 20, 30, and 40 m/s from the inside. The used data sets were recorded at meteorological stations in Aburatsu (Nichinan city, Miyazaki Pref.), Naha (Okinawa Island), and Ishigaki (Ishigaki Island). These meteorological stations were nearest to where the butterfly population dynamics were gathered.

Subsequently, geographic heat maps of the Ryukyu Islands and surrounding areas were generated to infer the impacts of typhoon disturbance based on their trajectories (Fig. 3c), and their association with the occurrence pattern in each year was assessed. Although there are variations yearly, these heat maps indicate that the more southerly areas are more affected by typhoons. This result is consistent with wind speed data (Fig. 4c), which indicates stronger winds in the southern regions during the typhoon season, lending reliability to the heat map. The estimated typhoon impact patterns for 2012 and 2013 exhibited similar trends; a strong influence in the more southerly part and the typhoons approaching Ishigaki Island with tremendous intensity. The occurrence patterns of these two years constituted cluster #2 for Ishigaki and cluster #3 for Okinawa, suggesting that typhoon disturbance affects population dynamics. Additionally, the occurrence patterns for Ishigaki Island in 2011 and 2015, when typhoons had a less significant impact on the island, were similar to those of Okinawa and Miyazaki and thus were included in cluster #4. Despite experiencing relatively low typhoon disturbance in 2014 in Ishigaki, only 31 individuals were recorded throughout the year with minimal fluctuations, resulting in it being included in Cluster #1.

### Effects of environmental parameters on the population dynamics of *Hebomoia glaucippe*

Since typhoons appeared to impact the population dynamics of *H. glaucippe*, we conducted a more comprehensive examination of the relationship between specific environmental variables and population dynamics based on data from Ishigaki Island and Okinawa Island. We selected mean temperature, maximum wind speed, and precipitation as the environmental variables, as these three factors are crucial as developmental conditions for organisms and are strongly related to seasonal adaptations of insects and host plants in subtropical regions. We observed a positive correlation between precipitation and wind speed on both Okinawa Island and Ishigaki Island (Fig. S8a, d), probably due to typhoons imparting strong winds and copious rainfall simultaneously. Conversely, the temperature did not correlate with the other two environmental variables on either island (Fig. S8b,d,e,f). Ascertaining the yearly or monthly relationship between these three environmental variables and population size in the correlation diagram was challenging (Fig. S9). Therefore, we analyzed the causal relationship by employing Convergent Cross-Mapping (CCM) based on the premise that the accumulation of environmental signals over several days influences population dynamics after a certain period (For detailed methods and experimental conditions, see **Materials and Methods** and Fig. S10).

The results indicated a causal relationship between mean temperature and abundance on Okinawa Island, maximum wind speed and abundance, and precipitation and abundance on Ishigaki Island (Fig. 5, Table 1). This examination of the causal relationship between mean temperature and abundance on Okinawa Island revealed that the optimal combination of time lag (n_1_) and signal accumulation window (n_2_) for maximum prediction accuracy was (n_1_, n_2_) = (25, 58), although all combinations of n_1_ and n_2_ examined in this study showed causal relationship (Fig. 5, Table 1). The S-map coefficients, showing the effect of environmental signal accumulation on population fluctuations, were consistently negative or zero with the optimal parameters (Fig. 6d). However, this result was due to both the *H. glaucippe* occurrences and the average temperature on Okinawa Island exhibiting an annual periodicity with synchronized cycle (Fig. 6a). The effect of environmental accumulation varied from negative to positive when the time lag changed (Fig. 6g).

**Figure 5.**
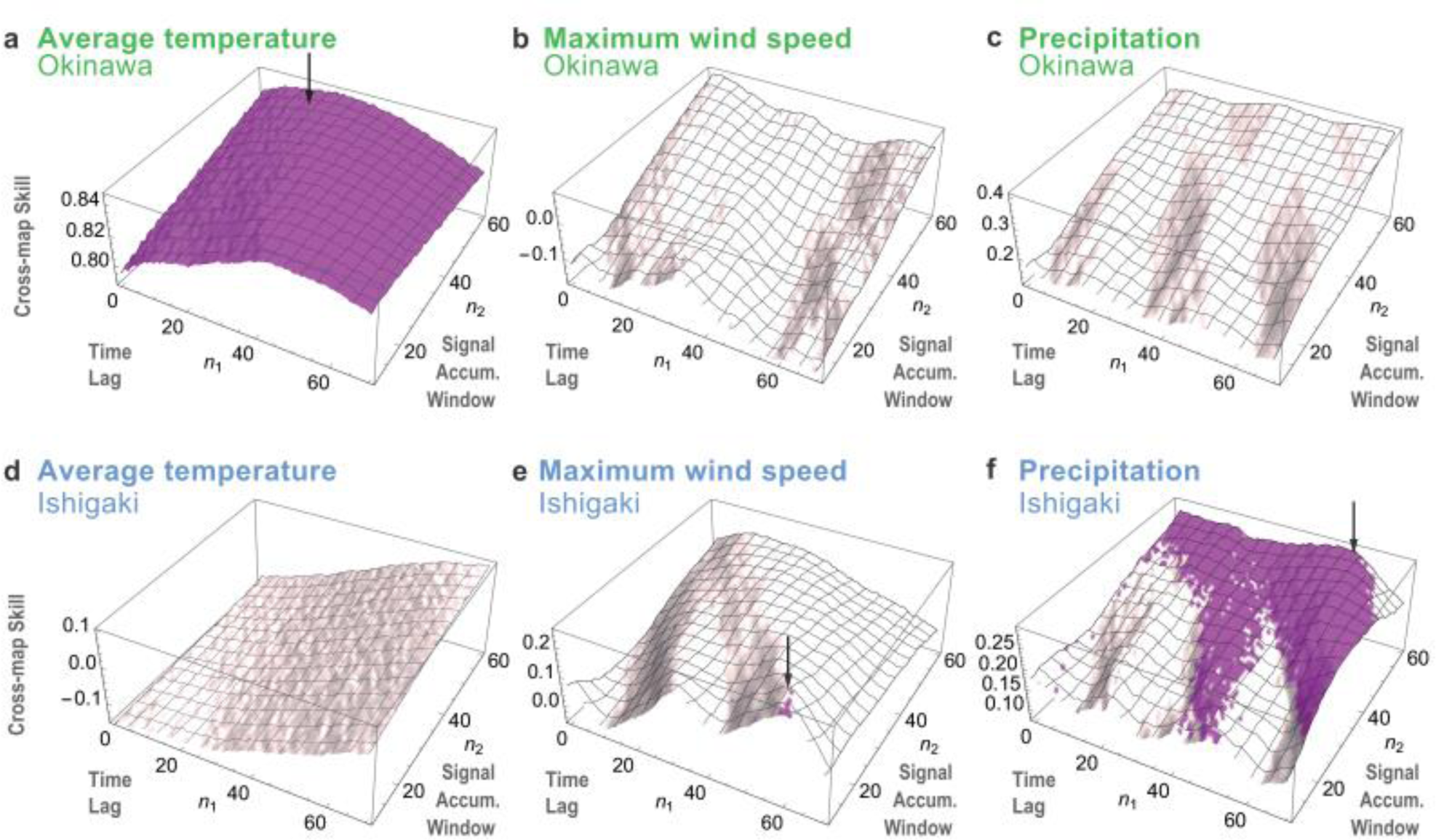
Causal relationships between population dynamics of *H. glaucippe* and environmental signals on each island. For the time series of *H. glaucippe*, a 30-day accumulation window was used here. The cumulative environmental signal was calculated for each date within the interval of cumulation (30 days) of *H. glaucippe* and averaged. Cross-map skill with maximum library size (⍴_max_) of the population dynamics in Okinawa Island **(a-c)** and Ishigaki Island **(d-f)** cross-mapping cumulative sum of average temperature **(a, d)**, maximum wind speed **(b, e)**, and precipitation**(c, f)**. Violet and white parameter regions indicate the causal and noncausal relationship, respectively. Arrows indicate the optimal parameter sets with the causality that provide the highest cross-map skills.

**Figure 6.**
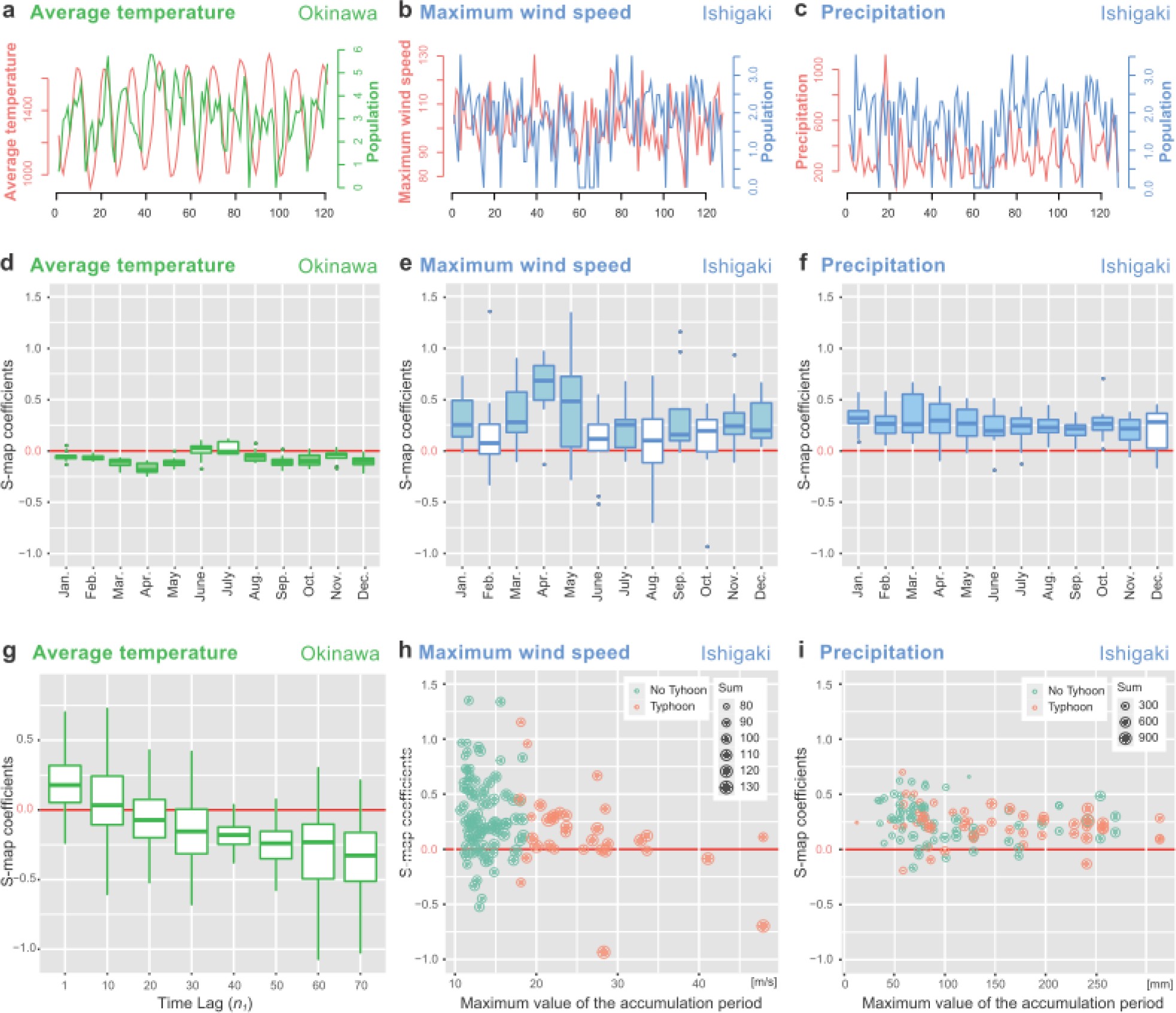
Effect of each environmental signal on the *H. glaucippe* population dynamics. **(a-c)** Projections of the population dynamics (the 30-day cumulative sum) and optimized environmental signals in the optimal parameter sets for which causality was detected. **(d-f)** Boxplots of estimated S-map coefficients for each month. Filled boxes indicate that the lower or upper 95% confidence intervals (CI) of median values are greater than or less than zero, respectively, indicating a significant difference from zero. **(g)** Boxplot of estimated S-map coefficients for the varying time lag (n1) conditions. **(h, i)** Scatterplot of estimated S-map coefficients and maximum values for environmental parameters. The size of the dots represents the total values of environmental factors during the time periods. The optimal parameter sets (n_1_, n_2_) for calculating the environmental time series in the Okinawa population with the average temperature **(a, d)**, the Ishigaki population with the maximum wind speed **(b, e, h)**, and with the precipitation **(c, f, i)** were (25, 58), (60, 12), and (55, 57), respectively.

**Table 1.**
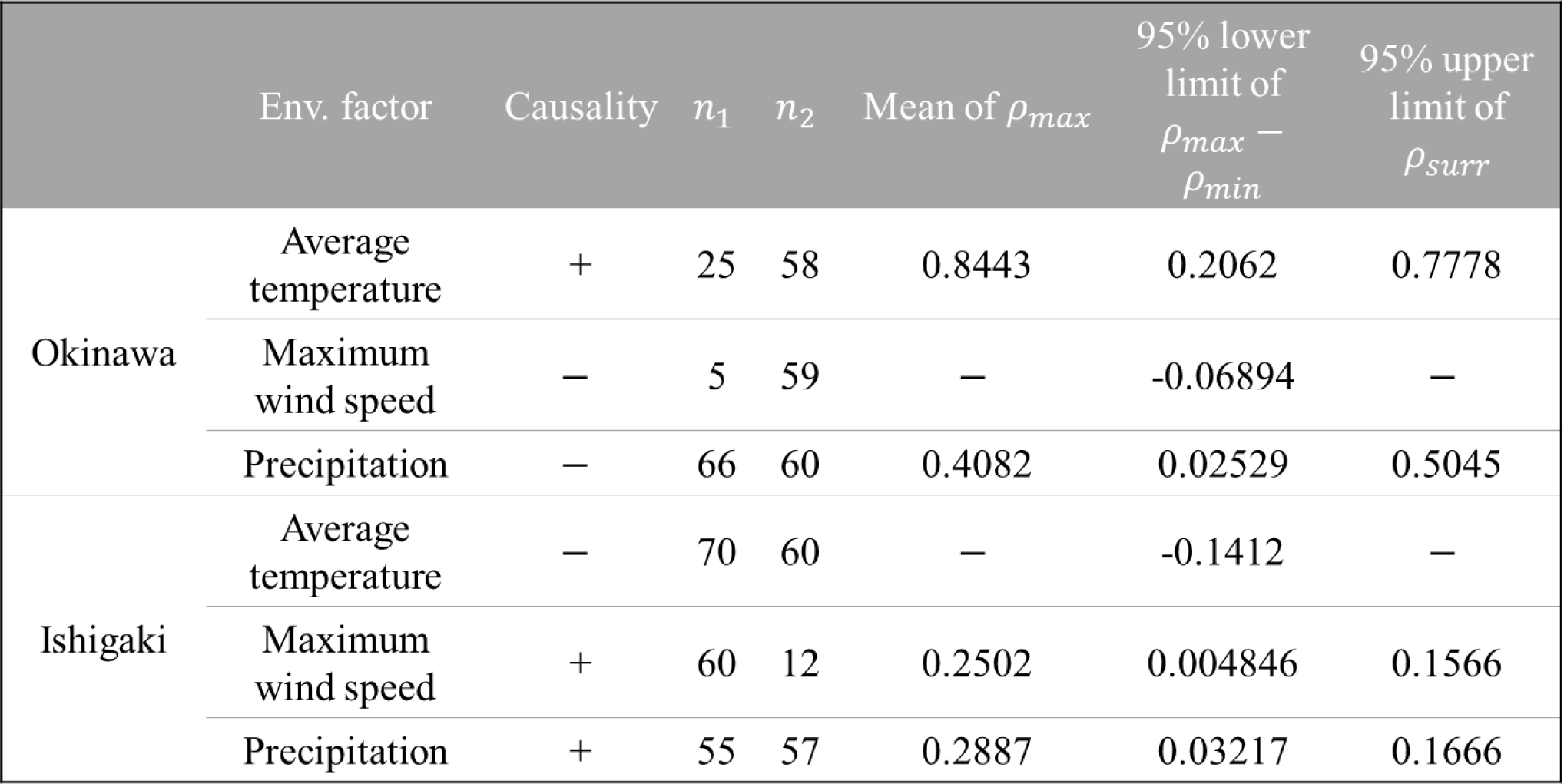
The convergent cross-mapping with the optimal parameter set (n_1_, n_2_). The following two criteria were used for identifying causality. First, the 95 % lower confidence limit of ⍴_*max*_ − ⍴_*min*_ is greater than null. Second, the mean of ⍴_*max*_ is larger than the 95% upper confidence limit of ⍴_*surr*_.

| | Env. factor | Causality | $n_1$ | $n_2$ | Mean of $\rho_{max}$ | 95% lower<br>limit of<br>$\rho_{max} - \rho_{min}$ | 95% upper<br>limit of<br>$\rho_{surr}$ |
| --- | --- | --- | --- | --- | --- | --- | --- |
| Okinawa | Average temperature | + | 25 | 58 | 0.8443 | 0.2062 | 0.7778 |
|  | Maximum wind speed | — | 5 | 59 | — | -0.06894 | — |
|  | Precipitation | — | 66 | 60 | 0.4082 | 0.02529 | 0.5045 |
| Ishigaki | Average temperature | — | 70 | 60 | — | -0.1412 | — |
|  | Maximum wind speed | + | 60 | 12 | 0.2502 | 0.004846 | 0.1566 |
|  | Precipitation | + | 55 | 57 | 0.2887 | 0.03217 | 0.1666 |

On Ishigaki Island, the maximum wind speed and population dynamics were found to be causal only for a few combinations around (n_1_, n_2_) = (60, 12), which is the optimal parameter set (Fig. 5e). This implies that the wind during the 12 days affected the population after 60 days, as the time lag of 60 days roughly corresponds to the time from egg to adult in *H. glaucippe* (Fig. 1b, c). Although the accumulated maximum wind speed had an almost positive effect on the population throughout the year, the S-map coefficients exhibited high variability and showed notably negative values during the typhoon season (Fig. 6e). More evaluation of the relationship between the maximum wind speed and the S-map coefficients indicated that a larger maximum wind speed could have a negative effect on the population in a few cases rather than the total maximum wind speeds for the 12 days (Fig. 6h).

Regarding precipitation and population on Ishigaki Island, a causal relationship was observed for time lags around 40 or 60 days, with the optimal parameter set being (n_1_, n_2_) = (55, 57) (Fig. 5f). The accumulated precipitation had a nearly uniform positive impact on the population throughout the year, and the S-map coefficients were stable, regardless of typhoon approach and maximum precipitation during the accumulation period (Fig. 6f, i). We found that the optimal time lags (n_1_) for the maximum wind speed and precipitation are 60 and 55 days, respectively, closely aligning with the generation time of *H. glaucippe cincia* (Fig. 1b,c). This indicates that both environmental factors impact the next generation, despite the differing signal accumulation windows (n2) and S-map scores.

## Discussion

Our study demonstrated that the three subspecies of *H. glaucippe* exhibit distinct overwintering strategies and that non-dormant individuals utilize Lammas growth induced by typhoon disturbance in the Yaeyama Islands. Furthermore, we analyzed the causal relationship between environmental parameters and butterfly population dynamics and found that non-dormant populations are influenced by typhoon-related environmental parameters, such as wind speed and precipitation. Collectively, these results suggest that typhoon-induced environmental disturbances, as well as seasonal climate change, significantly impact the wintering strategies of *H. glaucippe* in the Ryukyu Archipelago. While previous studies have discussed the impact of a single or a few typhoons on insects in a particular habitat or individual stray butterflies (e.g., Yukawa et al., 2020; Ogawa, 2022), our analyses enable us to elucidate how typhoons affect the population dynamics of insects based on more reliable meteorological evidence along with their geographic distributions.

Our field and rearing observations indicated that *H. glaucippe* populations exhibit varying degrees of dormancy within their life cycle. Similar dormancy variation by latitude has been reported in other Pieridae butterfly species in the Ryukyu Islands (Hashimoto et al., 2008; Yata, 1979). However, dormancy within the same habitat is consistent, contrasting with the observed variation in ssp. *liukiuensis* on Okinawa Island. Since our rearing experiments did not consider and restrict the genetic background of the derived from the same habitat, it is uncertain whether the variation in dormancy of ssp. *liukiuensis* is genetically determined or influenced by other subtle environmental factors that might modulate the threshold of diapause induction. Nonetheless, we observed that the pupal period of dormant individuals is about 75 days in ssp. *liukiuensis*, which is considerably shorter than that of approximately 120 days in ssp. *shirozui*, indicating that the dormancy trait of each subspecies has been adapted to suit the winter length of its habitat.

Alternatively, the non-dormant life-cycle may be considered an extension of the host adaptation in the nominotypical subspecies in Hong Kong (Bascombe et al., 1999). The larvae of *H. glaucippe* demonstrate remarkable cold tolerance, surviving temperatures below freezing (Fig. S4). Although they cannot molt under cold conditions (Fig. 1f), temperatures in the southern region of the Ryukyu Islands rarely drop to a level that inhibits larval molting (Fig. 4a). This suggests that the dormancy of *H. glaucippe* in Japan may not solely be attributed to low temperatures, but rather may be an adaptation to the host plant, allowing the larvae to overwinter without dormancy as long as a food source is available. However, predicting the food source availability remains challenging due to the unpredictable timing, size, and frequency of typhoons in the Ryukyu Islands (Fig. 3c). Thus, the seasonal phenology changes of *Crateva religiosa* are obscured by Lammas growth induction caused by typhoon disturbance, potentially counteracting the seasonal response for *H. glaucippe*. Consequently, the selective pressure for overwintering with dormancy may have weakened, leading to dormancy loss in ssp. *cincia* in the Yaeyama Islands. In essence, typhoons and the resulting changes in host plant availability may significantly shape the life-cycle and adaptative strategies of *H. glaucippe*.

The outcomes of our time-series analysis conducted on Okinawa and Ishigaki Islands populations are broadly consistent with our captive experiments and field observations. The causal relationship between population dynamics and temperatures on Okinawa Island was strong, with a high seasonal periodicity, as shown by CCM analysis (Fig. 5a, 6a). This finding suggests that during the winter months, food availability is limited, and the dominant over-wintering mode is through dormant pupae (Fig. 5b). In fact, we have observed dormant pupae in the field on Okinawa Island, indicating that the butterflies can overwinter as a larva only in years when environmental conditions are favorable. A significant causal relationship was observed between population size and both wind speed and precipitation in the Ishigaki population (Fig. 5e, f). The observed positive correlation between precipitation and wind speed was attributed to typhoons (Fig. S8). The optimal parameters for wind speed were (n_1_, n_2_) = (60, 12), while for precipitation, they were (n_1_, n_2_) = (55, 57). Although n_1_, the time lag, was similar, the value of n_2_, the integration period, differed considerably. This time lag is close to the generation time of *H. glaucippe cincia* (Fig. 1b,c), indicating that both environmental factors affect the next generation. The difference in the accumulation period may reflect how wind and rain influence *C. religiosa*. Strong winds caused by typhoons drop leaves and induce the Lammas growth, which may thereafter positively impact the population of *H. glaucippe*. However, they also have an immediate negative effect on butterfly development. Larvae residing on host leaves during typhoon strikes would be blown away with the leaves, and strong winds would directly threaten the survival of adults. In cases of consecutive typhoons, the negative effect would be more strongly observed if *C. religiosa* had already been defoliated by the previous typhoon. In fact, the S-map values of wind speed varied widely and were sometimes significantly negative when the maximum value of the integration period was high (Fig. 6h). This could be attributed to both positive and negative effects on the butterfly population. On the other hand, the S-map values of precipitation were stable throughout the year, which could be interpreted as a positive effect of precipitation on the population of *H. glaucippe* by stimulating the growth of *C. religiosa*.

In the present investigation, we have demonstrated that *H. glaucippe* exhibits a fascinating dormancy pattern, yet this species holds immense potential as a promising target for dormancy and overwintering strategies. The lack of discernible morphological disparities between dormant and non-dormant pupae of *H. glaucippe* implies that dormancy may not have fully evolved in this species (Yata, 1977). Thus, it would be advantageous to investigate the early stages of dormancy acquisition. Furthermore, in Taiwan, ssp. *formosana* Fruhstorfer, 1908, which belongs to the *glaucippe* group represented by the nominotypical subspecies, is found across a range of elevations and habitats (Morishita, 1973; Hamano, 1986; Samusawa, 2021). Additionally, *H. glaucippe* utilizes more than seven species of Capparaceae as hosts, including evergreen species (Hong, 2020), indicating that *H. glaucippe* may have undergone unique life history evolution in Taiwan compared to the Ryukyu Islands. An in-depth comprehension of the ecological and genetic foundation of this species could provide additional insights into these evolutionary processes.

A noteworthy aspect of our study is the innovative integration of citizen science to collect population dynamics data. With the global presence of insect enthusiasts, a vast reservoir of ecologically valuable data remains untouched. We capitalized on such data by employing advanced statistical techniques for analysis. Although our study focused on the impact of typhoon-induced disturbances, retrospective data collection before a specific event can pose challenges. However, in situations where unpredictable environmental events are the focus, the benefits of assembling and utilizing pre-existing data are substantial. Furthermore, we implemented the Convergent Cross Mapping (CCM) method, a tool of Empirical Dynamic Modeling (EDM), to establish causal relationships between variables in conjunction with basic statistical analysis. This methodology has recently gained traction in ecological research (Kawatsu & Kishi, 2018; Ushio et al., 2018; Ueno et al., 2021; Satake et al., 2021). Our mathematical analysis conducted on populations of Okinawa and Ishigaki Islands is broadly consistent with our captive experiments and field observations, thus indicating the effectiveness of our approach for ecological studies that focus on the influence of unpredictable and unstable environmental conditions, such as hurricanes, river flooding, and cliff collapses.

## Materials & Methods

### Field Observations

From 2017 to 2022, field observations of *Hebomoia glaucippe* were conducted in Miyazaki Prefecture (Nichinan City), Kagoshima Prefecture (Ibusuki City), Yakushima Island, Amami-Oshima Island, Okinawa Island, Miyako Islands, Ishigaki Island, Iriomote Island, and Yonaguni Island. One of the co-authors, F. Ishiwata, resided on Iriomote Island from April 2019 to March 2021, Miyazaki Prefecture from April 2021 to March 2022, and Yakushima Island from April 2022, conducting year-round field observations in each location. We also conducted multiple surveys in regions other than these, mainly during the overwintering season. During the field surveys, we looked for the sacred garlic pear *Crateva religiosa* tree, the host plant of *H. glaucippe*, and noted the presence of *H. glaucippe* and the condition of *C. religiosa*.

### Rearing Experiment of the three subspecies of *H. glaucippe*

A rearing experiment was conducted to compare the diapause and cold tolerance of *H. glaucippe* in each region. In the rearing trials, eggs collected in the field or ones deposited by adult females in captivity were employed. Throughout the indoor raising experiment to evaluate diapause duration, the room environment was held at 12L12D 20°C (pupae of this species do not require cold stimulation to emerge, so the pupae were maintained under the same conditions). The larvae were maintained in 220 mm x 150 mm x 160 mm mesh-topped plastic cups containing leaves of *C. religiosa* inserted into a small water pot., with about five larvae per container up to the third instar and one larva per container from the fourth (subterminal) instar. Larval and pupal periods (days) for each subspecies were recorded and compared using one-way ANOVA followed by a post-hoc Tukey HSD test. RStudio (version 2021.09.0, Build 351) was used for the statistical analysis. In the field breeding experiment to confirm whether the butterfly can withstand severe winter conditions, transplanted *C. religiosa* trees on the Kyushu University campus in Fukuoka Prefecture, which is north of the northern border of *H. glaucippe* distribution, were employed. When larvae died during the experiment, the cause of death was also noted.

### Collection of Long-term Population Dynamics data and Meteorological data

The number of adult butterfly counts was employed as data showing the population dynamics of *H. glaucippe*. These data were extracted from journals published for insect enthusiasts in Japan (see Supplemental Table S1 for detail). For subspecies *shirozui*, occurrence data were available for the most northern population in Miyazaki prefecture. This dataset contains 286 individual records for the four years from 2015 to 2018, averaged for the early-mid-late of each month. For subspecies *liukiuensis* and *cincia*, daily data were available for 5705 and 894 individuals recorded from January 2010 to December 2019 on Okinawa Island and from August 2009 to February 2020 on Ishigaki Island, respectively.

Regarding meteorological data, daily records of average temperature, precipitation, and maximum wind speed were downloaded from the Japan Meteorological Agency’s database (https://www.data.jma.go.jp/obd/stats/etrn/index.php). We used datasets from Aburatsu (Miyazaki), Naha (Okinawa Island), and Ishigaki since these observation points are closest to where the population data were recorded. The number of approaching typhoons (defied as it entered within 300 km of a meteorological observing station) was also obtained from the Japan Meteorological Agency’s database. The typhoon tracking data was collected from the Joint Typhoon Warning Center’s best tracking database (https://www.metoc.navy.mil/jtwc/jtwc.html?western-pacific), pinpointing the location of each typhoon at 6-hour intervals.

### Visualization and Statistical analysis of the datasets

Long-term population dynamics data, temperature, and approaching typhoons were visualized as heat maps using the ‘superheat’ package (version 0.1.0) in RStudio (version 2021.09.0). The superheat package’s scaling and clustering options were employed to standardize population data for comparing emergence patterns among years and regions and hierarchically clustered with the ward method. The clusters were identified using dendrogram shape and silhouette analysis, with a maximum average silhouette value observed at four. Temperature and approaching typhoon heatmaps were created without scaling to compare regional differences. Percentile roses to visualize the daily maximum wind speed and wind direction were created using the ‘openair’ package (ver. 2.8.6). The percentile was calculated based on the daily maximum wind speed for each location. Heatmaps for estimating areas of typhoon disturbance were created using QGIS (ver. 3.26) by mapping the center location of the typhoon every six hours. The ACF analyses were conducted using the “acf” function from the “stats” package (version 3.6.3) in R software (version 3.6.3). These analyses were applied to the daily time-series data, examining time lags ranging from 1 to 400. Correlation coefficients were calculated using the “cor” function from the same “stats” package, with both Pearson and Spearman coefficients calculated. Additionally, a linear regression line was drawn using the “stat_smooth()” function from the “ggplot2” package (version 3.3.3).

### Cumulative time series used for causality test

The time series datasets were converted to cumulative values as time series variables for the causality test to reduce the influence of noise. The cumulative time series of *H. glaucippe* counting data was created as follows. The cumulative sum for 30 days was calculated, added ‘1’ to each time point, and log-transformed with the natural logarithm. Since *H. glaucippe* adult life span was estimated to be over one month (Iwasaki, 2019), a cumulative period of 30 days was employed. We assumed that the environmental signals cumulate over n_2_ days from n_1_ days before the sample collection date; therefore, the cumulative time series of environmental signals were created as follows. First, the cumulative environmental signal was calculated for each date within the interval of cumulation (30 days) included in each point of the time series of *H. glaucippe*. Then, the mean value of the cumulative signals calculated within the interval was defined as the environmental time-series point corresponding to the time-series point of *H. glaucippe* (Fig. S10). The time lag between the environmental signal calculation start and the butterfly emergence (n_1_) and the signal accumulation window (n_2_) varied by integers from 1–70 and 11–60 days, respectively.

### Causality test using convergent cross mapping (CCM)

We used equation-free non-linear time series analysis called Empirical Dynamic Modeling (EDM) to test whether the population dynamics of *H. glaucippe* were derived by temperature, precipitation, or maximum wind speed. Convergent Cross Mapping (CCM), one of the tools of EDM, is used to detect causality between variables. Our CCM analysis was carried out using the ‘rEDM’ package (version 0. 7. 1) in R (version 3.6.3) in three steps according to the standard protocol of EDM (Chang et al., 2017): (i) The optimal embedding dimension *E* for the state space reconstruction of the time series of *H. glaucippe* was determined by simplex projection (Sugihara & May, 1990; Sugihara et al., 2012); (ii) The nonlinearity of the *E*-dimensional reconstructed attractor was checked by S-map (sequential locally weighted global linear map, Sugihara, 1994) and (iii) CCM was performed in the direction from the variable of *H. glaucippe* to environmental variables (temperature, precipitation, and maximum wind speed). We then determined the optimal set of (n_1_, n_2_) with the highest cross-map skill with respect to the environmental variables, which is causally related to the variable of *H. glaucippe*. We set time lag t = 1. Given that the maximum embedding dimension which can be used in the EDM framework is roughly the square root of the time series length of a given dataset (Munch et al., 2019), we searched for the optimal embedding dimension up to the integer less than or equal to the square root of the time series length. To test whether there is convergence, we calculated the cross-map skill with the minimum (= *E* + 1) and the maximum (= time series length – *E* + 1) library length using 1000 bootstrap samples (ρ_min_ and ρ_max_, respectively). In addition, to check the false high cross-map skill, we generated 100 seasonal surrogate data of the environmental signals, which preserve the seasonality of the original environmental time series but lose the causal relationship with the time series of *H. glaucippe*. Seasonal surrogates were generated by extracting a mean seasonal trend of the specified period (T-period) with a smoothing spline and shuffling the residuals. T-period was determined by dividing 365 by the signal accumulation window (= 30) of the time series of *H. glaucippe*. Then, we calculated the cross-map skill of the variable of *H. glaucippe* cross-mapping surrogate (*ρ*_surr_). We judged the causality significance using two criteria: (a) the 95% lower confidence limit of *ρ*_max_*–ρ*_min_ is greater than zero, and (b) the mean of *ρ*_max_ is larger than the 95% upper confidence limit of *ρ*_surr_.

### Measuring interaction strength from environmental signals to population dynamics of *H. glaucippe*

To estimate the interaction strength from the environmental signals to the population dynamics, we performed the multivariate S-map (sequential locally weighted global linear map, Chang et al., 2017; Sugihara, 1994) using the causal environmental signals at the optimal set of (n1, n2) determined by CCM. Furthermore, in the S-map analysis of butterfly occurrence and temperature in Okinawa, where causality was shown for all combinations of n1 and n2, the time lag (n1) was varied to assess the influence of the time series data’s high periodicity on the S-map. The S-map method calculates partial derivatives of affected variable X by affecting variable Y (i.e., Jacobian ∂X/∂Y (Deyle et al., 2016)) in a multivariate state space at each time point, and yields S-map coefficients that are good approximations of the interaction strength from Y to X at each time point. Replacing the last coordinate of the reconstructed *E* dimensional attractor of *H. glaucippe* with an environmental variable, we created a multivariate SSR attractor and carried out the multivariate S-map. We normalized each time series used for the S-map to have zero mean and unit variance, so Euclidian distance in the multivariate embedding is scale-free.

## Acknowledgment

We appreciate Mr. Daigo Enomoto and Dr. Ken-ichi Odagiri supporting our fieldwork and butterfly rearing. This research was funded by a JSPS KAKENHI Grant Number 21K15165 (Grant-in-Aid for Young Scientists) to KO. The fieldwork and mathematical analyses are partially supported by the Collaborative Research of Tropical Biosphere Research Center, University of the Ryukyus (Japan), and the Education and Research Center for Mathematical and Data Science, Kyushu University (Japan).

## Author contributions

KO and YM conceived the project. KO and FI carried out the field observation and rearing experiments. KO extracted data from the literature and performed statistical analyses. WN conducted and AS supervised the EDM (CCM) analysis. KO wrote the manuscript draft, and all authors read, commented on, and revised the manuscript.

