## Supplementary material for "Typhoon-induced Lammas growth promotes the non-dormant life-cycle of the Great Orange Tip butterfly *Hebomoia glaucippe*": Supplemantal Figures

**Supplemental Figures**

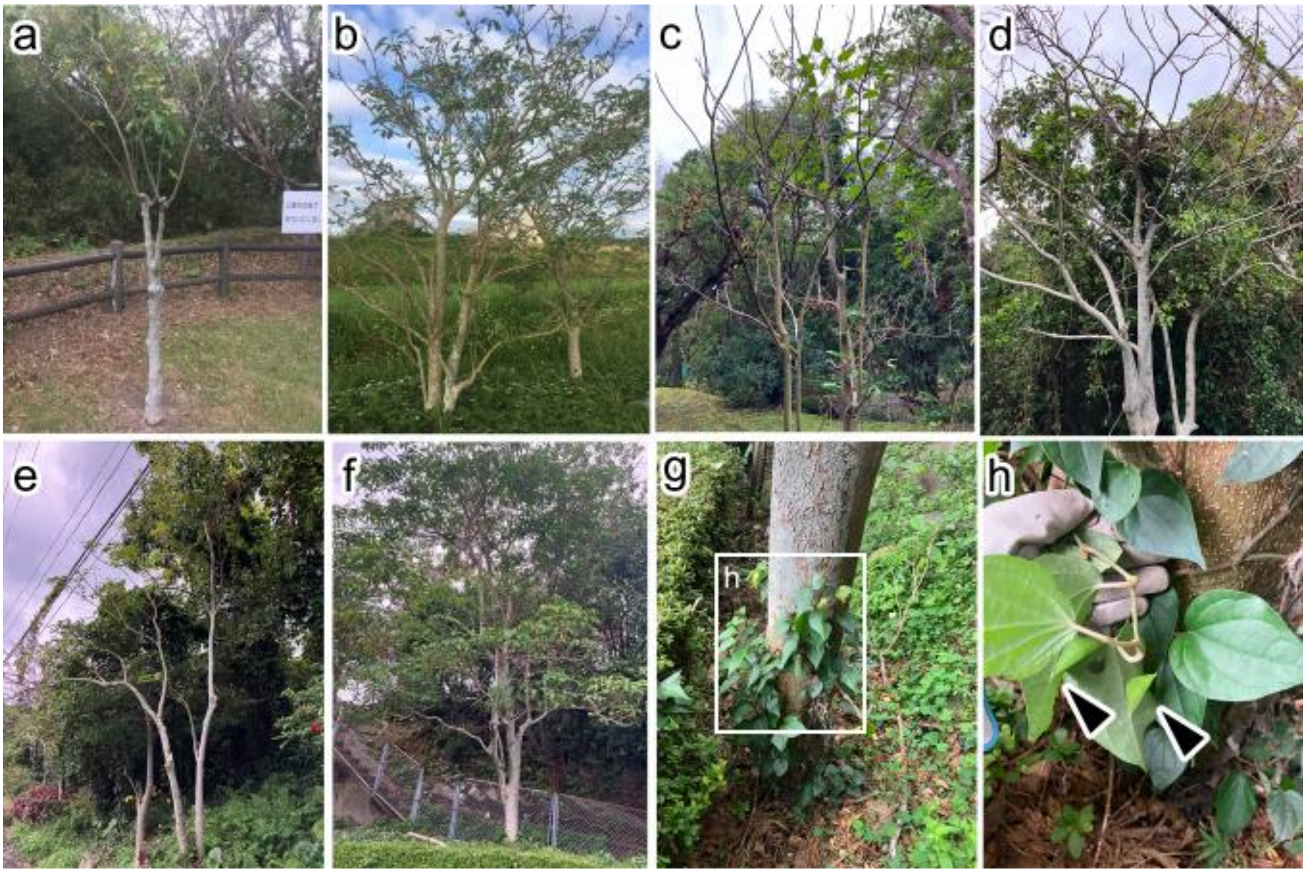


**Figure S1. The deciduous status of *Crateva religiosa* and the overwintering status of *Hebomoia glaucippe shirozui* in Kyushu (a, b) and Yakushima Island (c-h) during the autumn of 2022 to early spring of 2023.** Most *Crateva religiosa* trees on Kyushu and Yakushima Island exhibited deciduous characteristics during the winter season (a-f). Prior to complete defoliation, many pupae were found on the leaves and branches of plants nearby the host (g, h). These pupae were collected to analyze pupal duration under rearing conditions (12L12D 20°C). All individuals (14/14) that resulted in butterfly emergence had a pupal duration of 40 days or more, indicating they were dormant pupae.

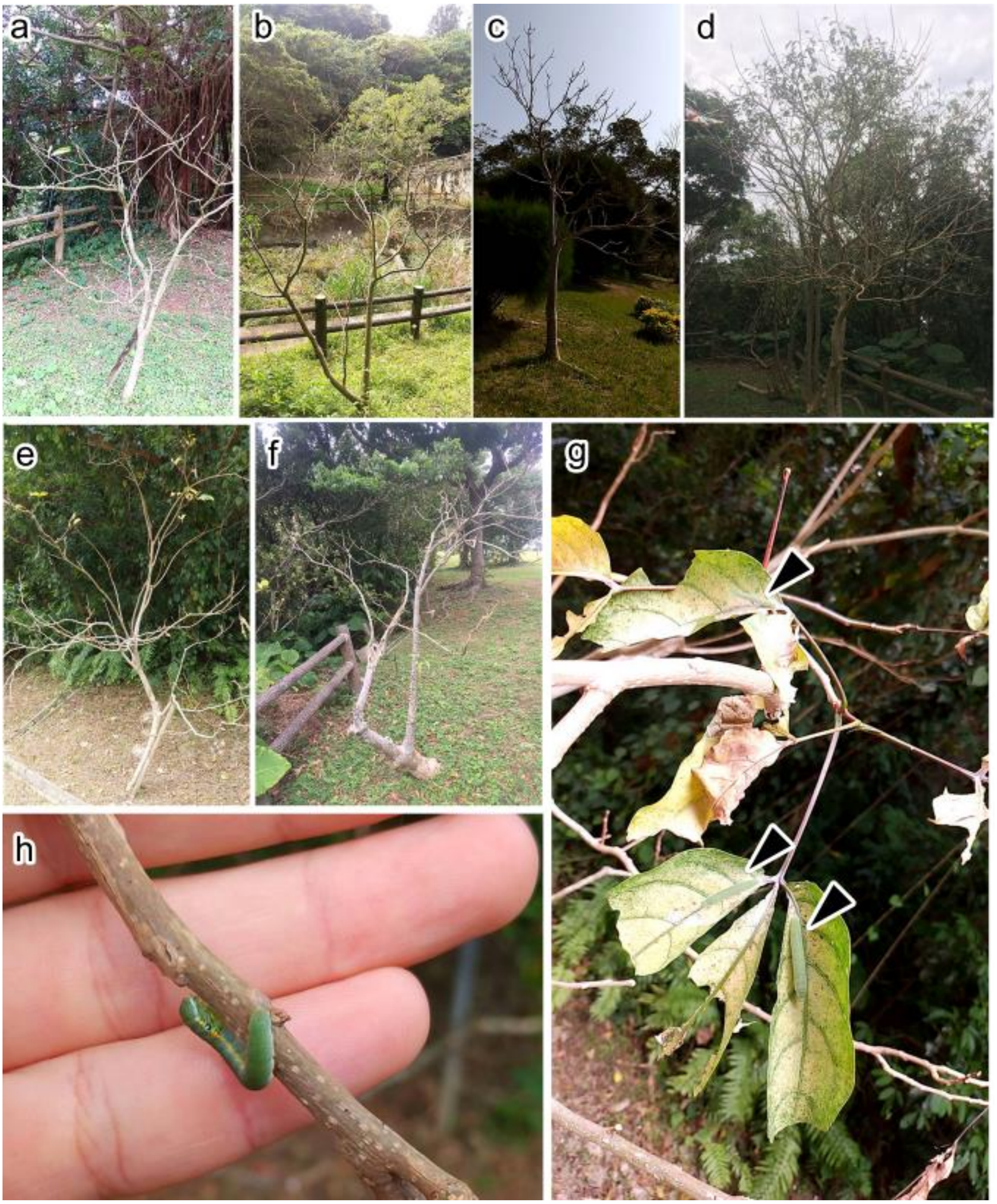


**Figure S2. The deciduous status of *Crateva religiosa* and the overwintering status of *Hebomoia glaucippe liukiuensis* in Okinawa Island during the autumn of 2019 to early spring of 2020.** By February 2020, defoliation was detected on most trees **(a-f)**. The few remaining leaves were too mature and close to falling off. On the remaining leaves, several larvae were found **(g)**. Some larvae were seen wandering the branches looking for leaves to feed on **(h)**. In 2019-2020, the success rate of larval overwintering in this region was deemed very low. Arrowheads indicate larvae on leaves. The images shown here were captured on February 27, 2020.

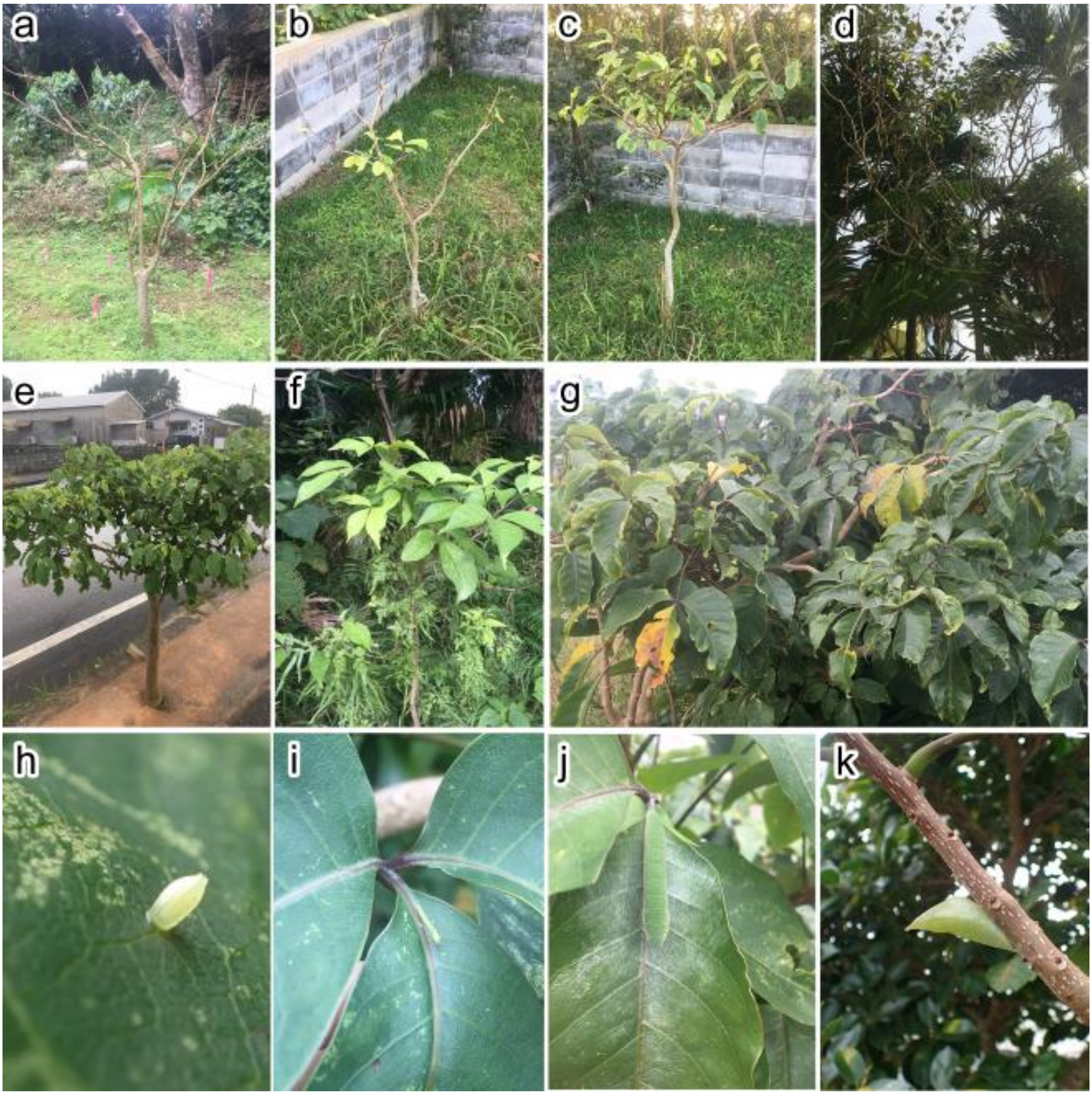


**Figure S3. The deciduous status of *Crateva religiosa* and the overwintering status of *Hebomoia glaucippe cincia* in Iriomote Island during the autumn of 2019 to early spring of 2020.** On the Yaeyama and Miyako Islands, including Iriomote Island, some *C. religiosa* trees lose their leaves during the winter, while others do not, indicating that the phenology is not synchronized even within the same island **(a-g)**. On the trees with healthy leaves, *H. glaucippe* eggs, larvae, and pupae were found **(h-k)**. The images shown here were captured on February 7-8, 2020.

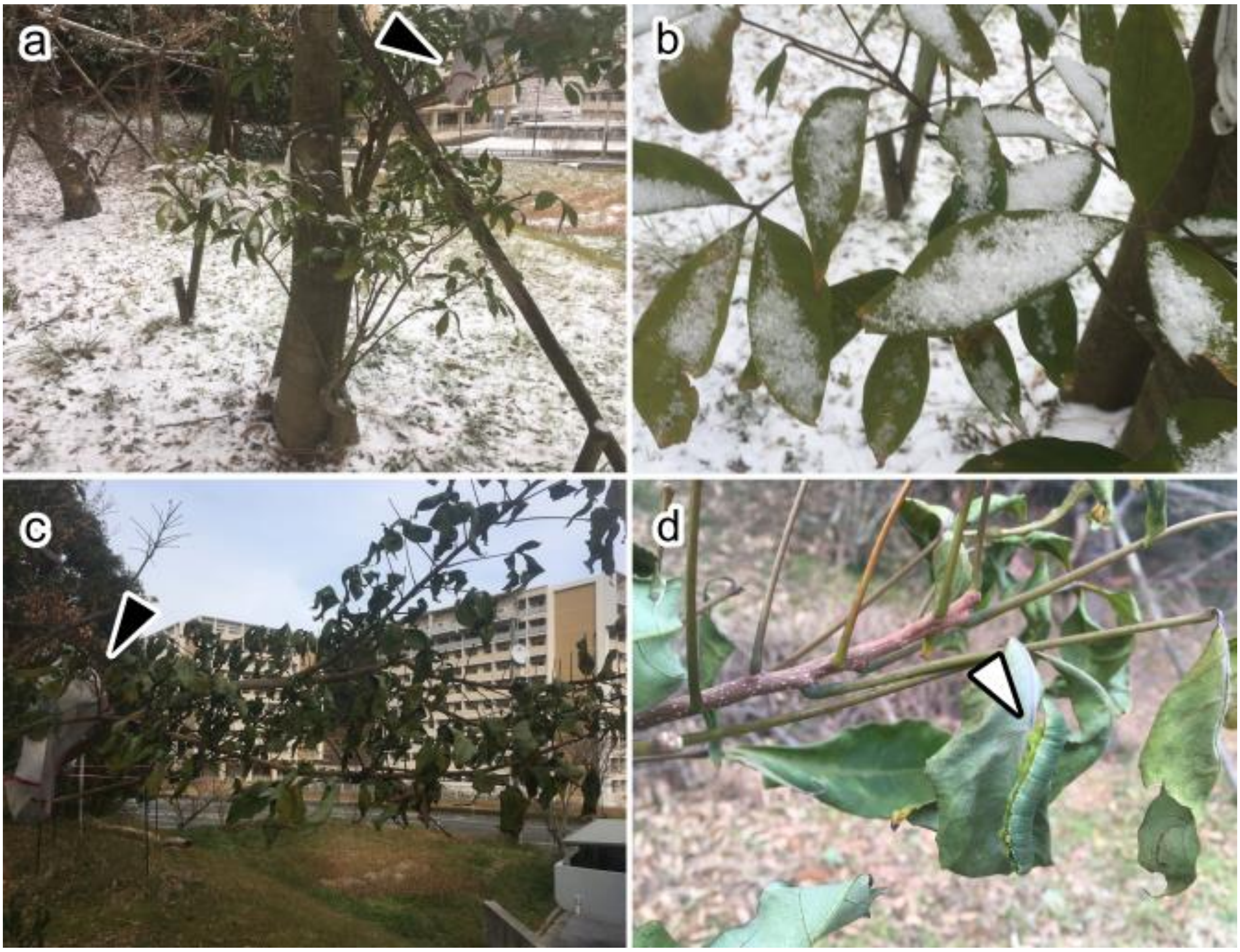


**Figure S4. Field rearing experiments in Fukuoka City and low-temperature tolerance of *Hebomoia glaucippe* larvae.** To evaluate the cold tolerance of *H. glaucippe* larvae, field rearings were undertaken in Fukuoka City (33°35'58N 130°13'22E), which is located farther north than their natural distribution. During the rearing experiment, a cold storm came, and snow accumulation was observed **(a, b)**, even though Fukuoka City's lowlands seldom get snowfall. Frostbite brought on by the snowfall and freezing temperatures caused the leaves to wilt **(c)**, but the larvae did not die **(d)**. Arrowheads in black and white indicate the rearing cage and the larvae on the wilted leaves, respectively.

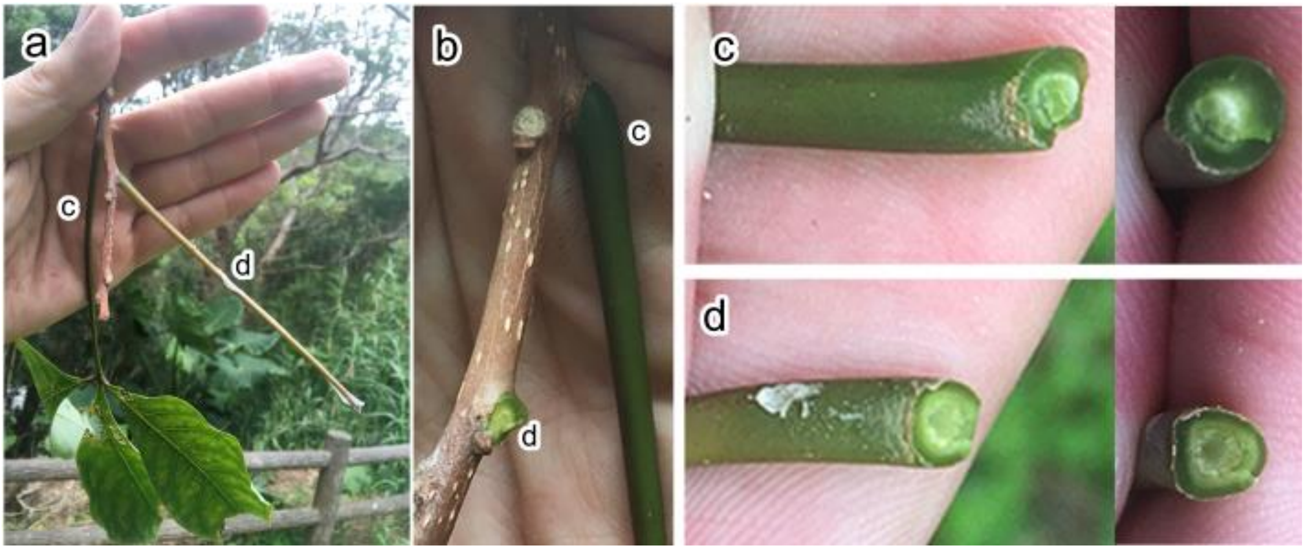


**Figure S5. Yellowing and defoliation occur slowly in *Crateva religiosa*.** The leaves of C. religiosa fall slowly, resulting in the presence of leaves at different stages on the same branch the following summer **(a)**. The petiole base separates from the branch when physical force is applied to the aged leaf **(b)**. The petiolar tissue stays alive even after the leaf blade is detached, and only the petiole remains **(c, d)**. These defoliation features may promote simultaneous defoliation triggered by typhoons and subsequent Lammas shoots sprouting.

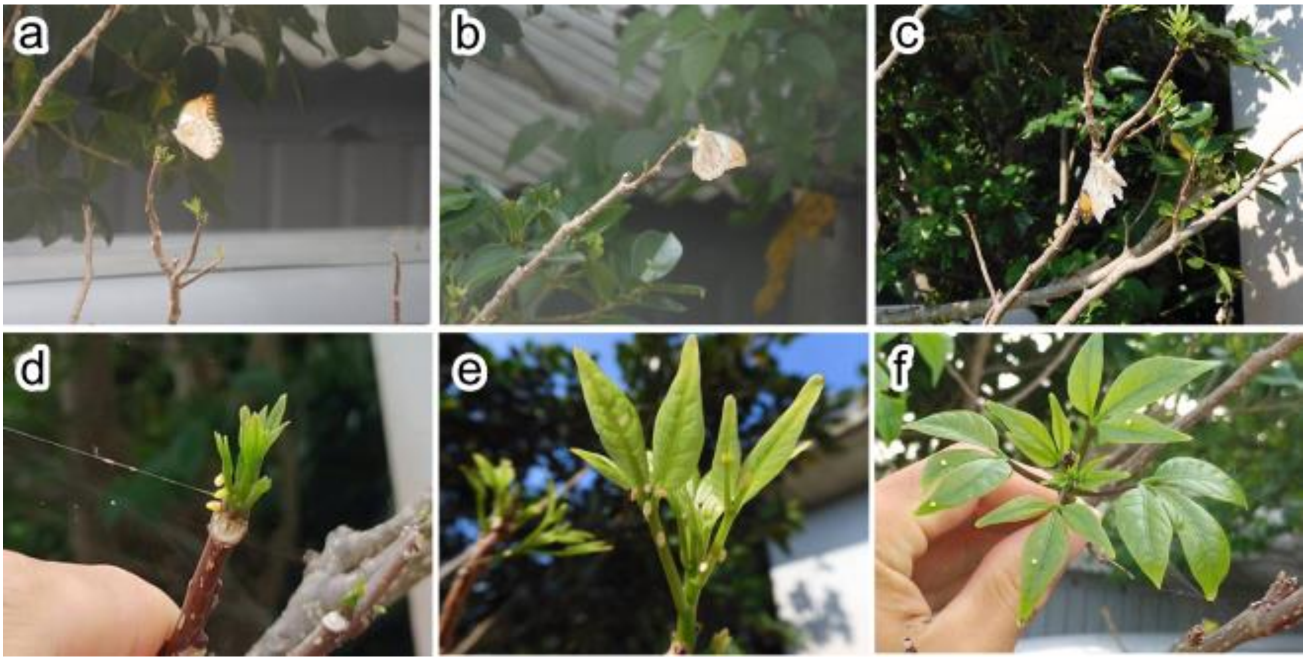


**Figure S6. Ferocious oviposition on the Lammas shoots. (a-c)** *H. glaucippe* ovipositing on the Lammas shoot. **(d-f)** Numerous eggs laid on the Lammas shoots. When the Lammas shoots are sprouted, *H. glaucippe* mother swarms to and lays eggs on them. The mother often lays eggs on the earliest shoots and young, slightly developed leaves but never on old leaves if there are sprouting shoots.

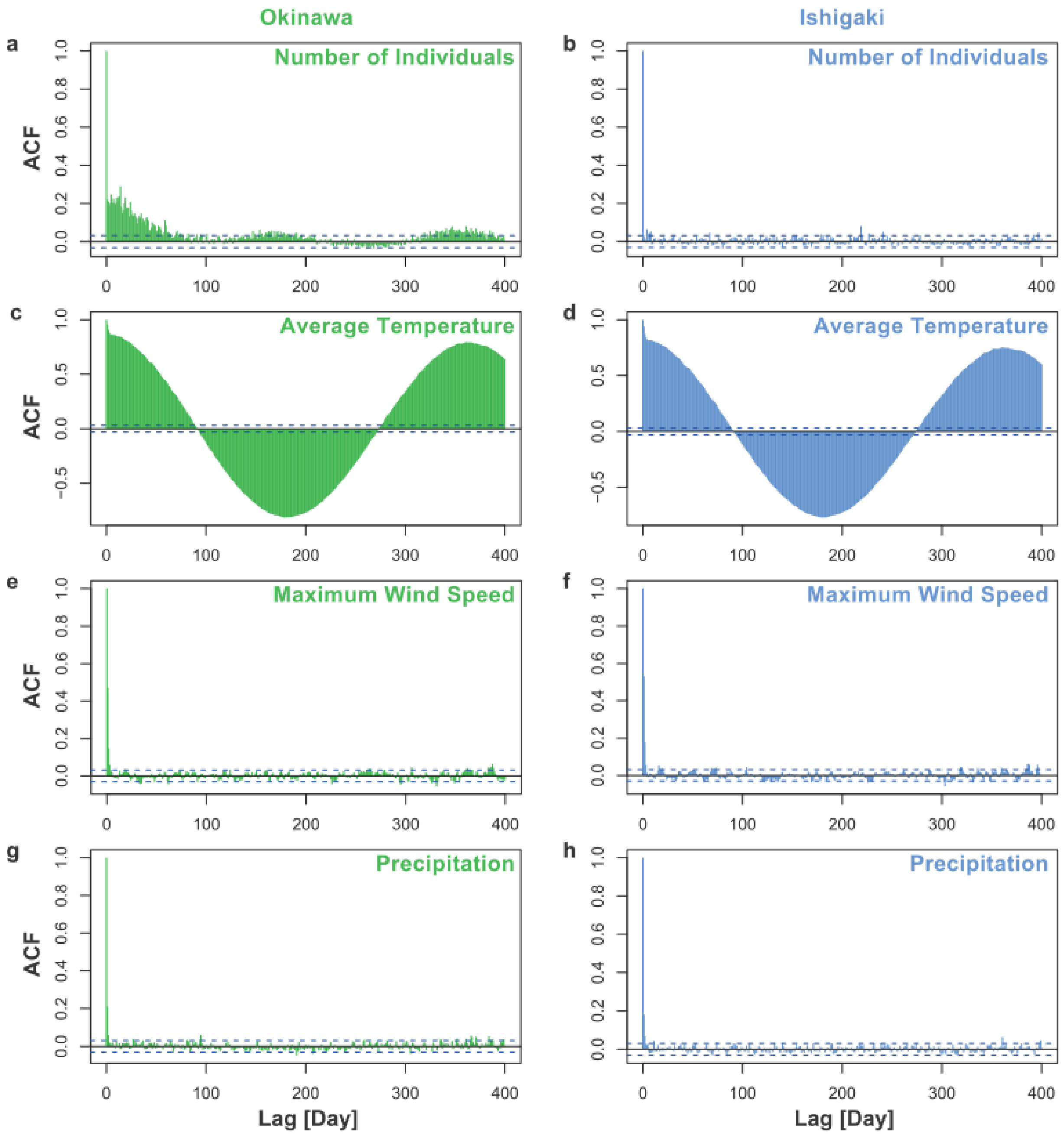


**Figure S7. Autocorrelation functions (ACF) analysis of time series data on the population dynamics and environmental parameters.** Histogram of the number of individuals **(a, b)**, the average temperature **(c, d)**, the maximum wind speed **(e, f)**, and the precipitation **(g, h)** in Okinawa island **(a, c, e, g)** and Ishigaki island **(b, d, f, h)**. The dashed lines represent the 95% confidence intervals for the null hypothesis that the autocorrelation values are zero for one or more lags.

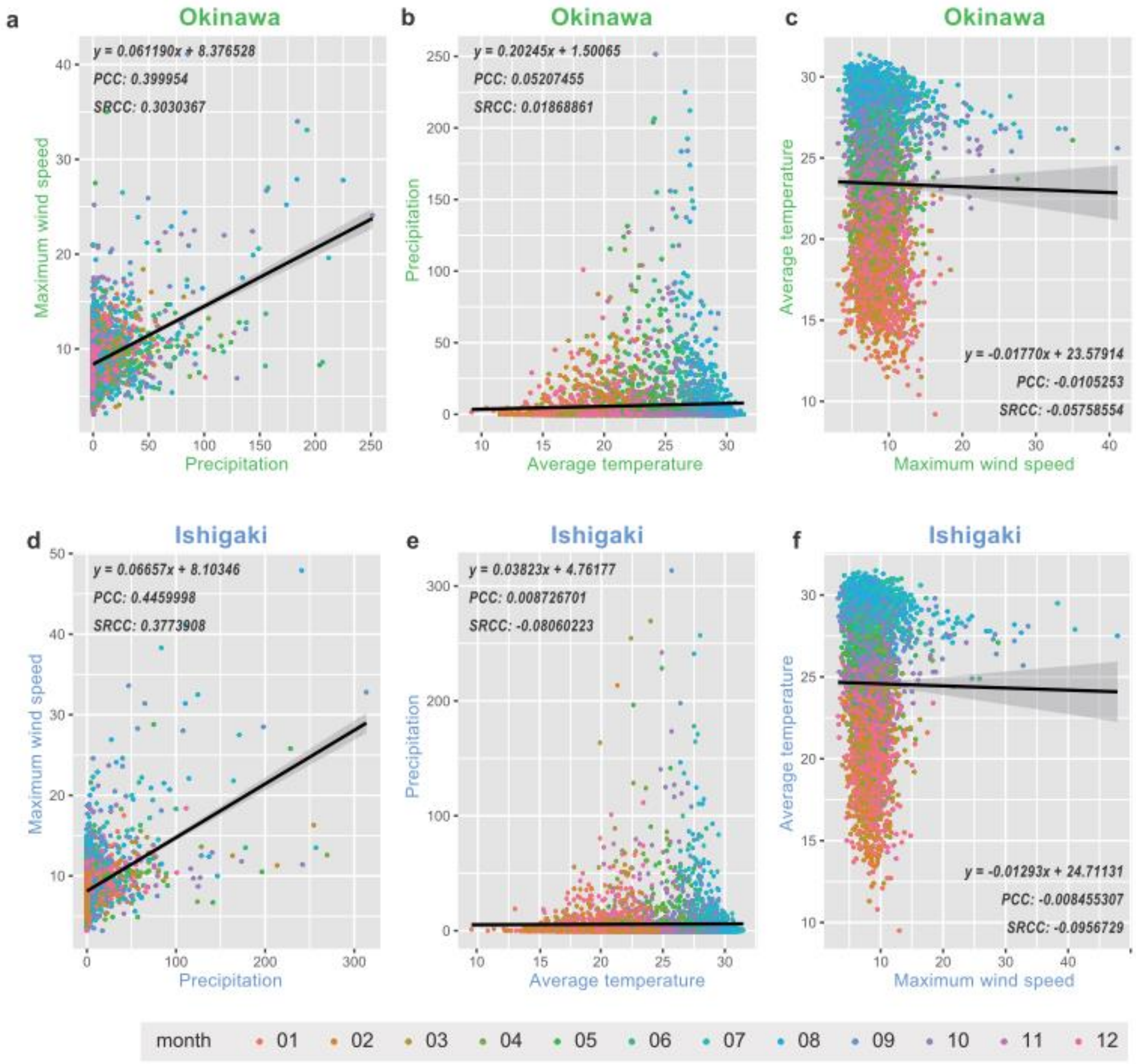


**Figure S8. The correlation analysis between the environmental parameters.** The scatter plots with the linear regression lines, Pearson correlation coefficient (PCC) and Spearman's rank correlation coefficient (SRCC) between the maximum wind speed and the precipitation **(a, d)**, the precipitation and the average temperature **(b, e)**, and the average temperature and the maximum wind speed **(c, f)** in Okinawa **(a-c)** and Ishigaki **(d-f)**.

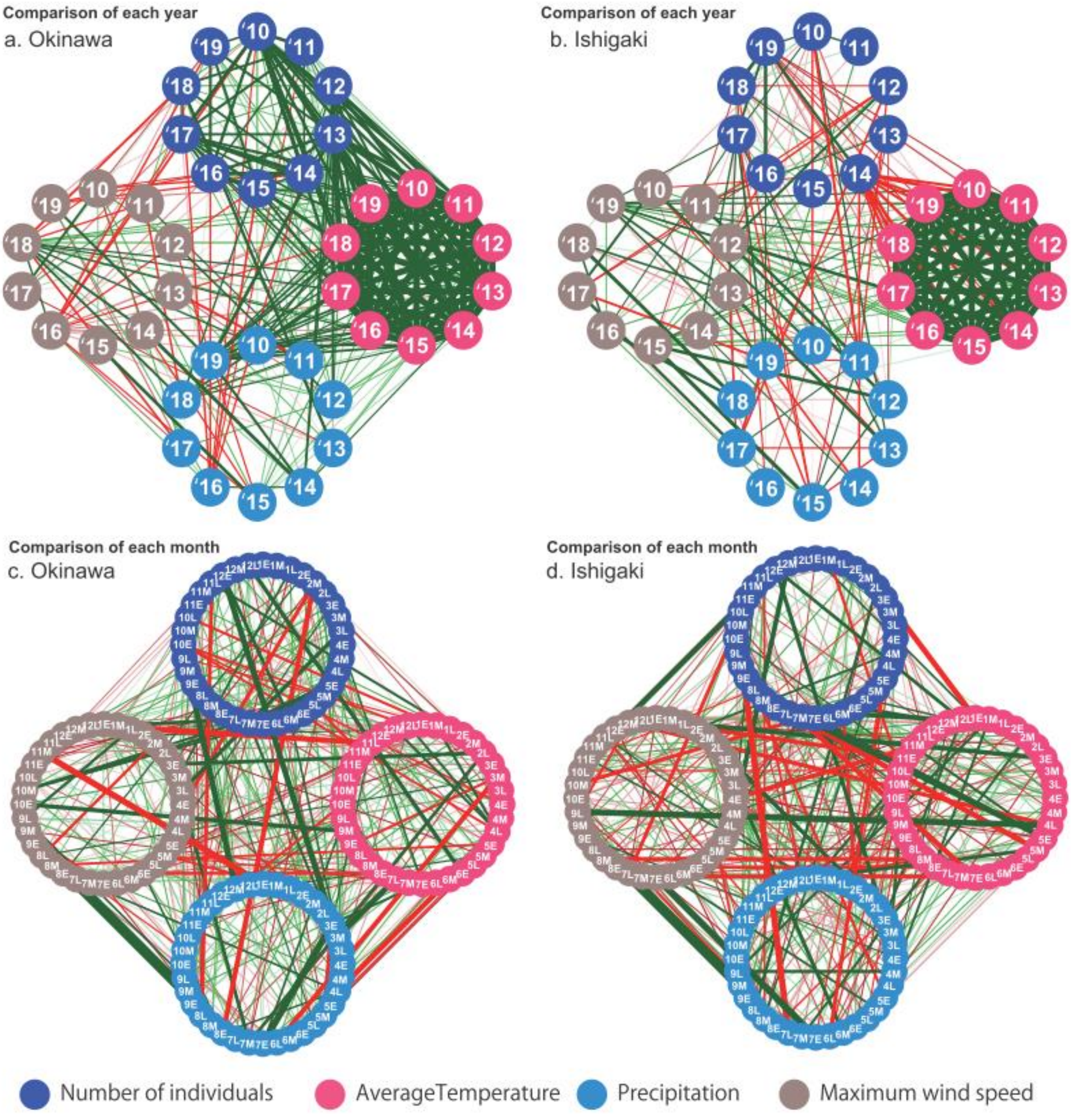


**Figure S9. Correlation diagram of the population dynamics and environmental parameters. (a, b)** Comparison for each year in Okinawa (a) and Ishigaki (b). **(c, d)** Comparison for each month divided into early-mid-late in Okinawa (c) and Ishigaki (d). The green and red lines indicate positive and negative correlations, respectively, with thicker lines indicating higher correlations.

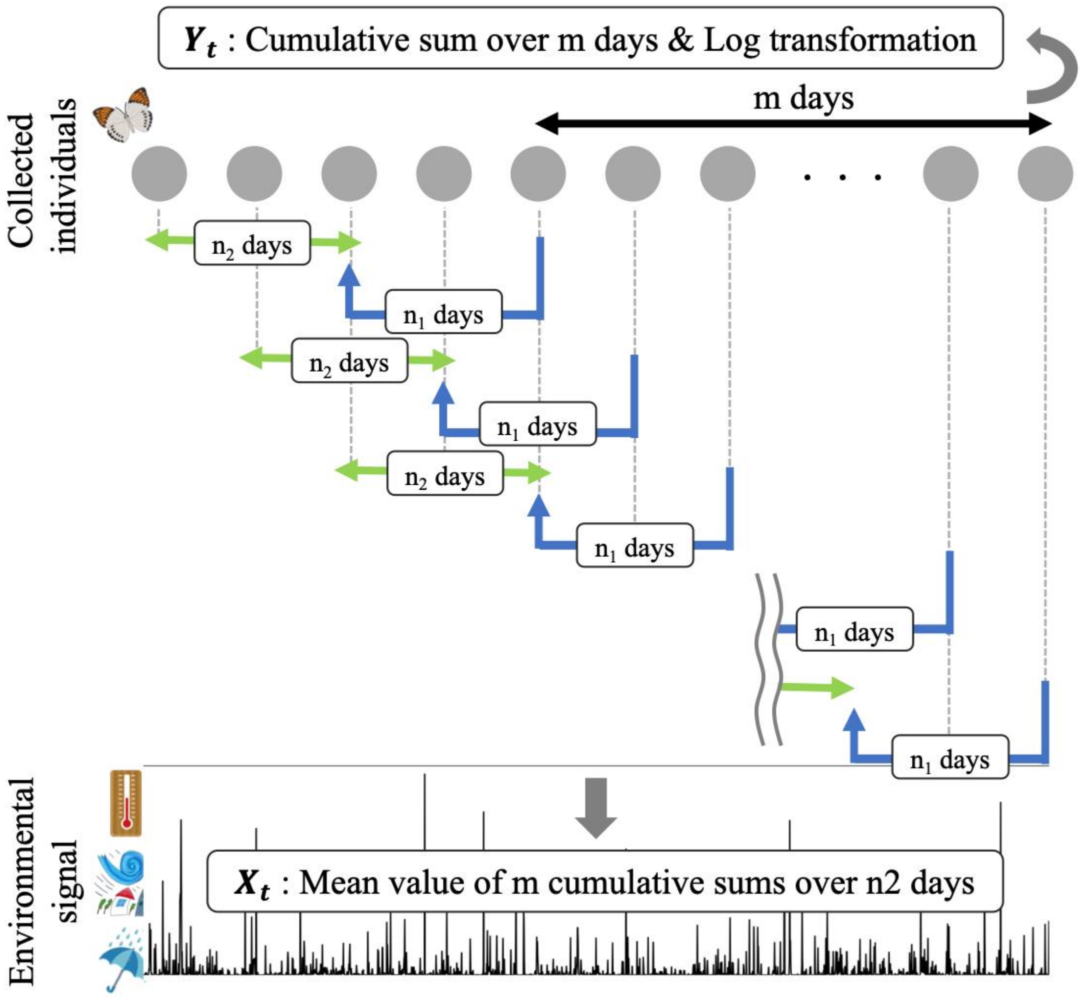


**Figure S10. Conceptual diagram of the CCM analysis**

**Supplemental Table**

**Table S1. List of references citing butterfly collection data** (All literature written in Japanese and translated into English by us)**.**
